# Capsid flexibility during Ty1 virus-like particle assembly

**DOI:** 10.64898/2025.12.17.694965

**Authors:** Bryan S. Sibert, J. Adam Hannon-Hatfield, Giuseppe Nicastro, Matthew A. Cottee, Ian A. Taylor, David J. Garfinkel, Elizabeth R. Wright

## Abstract

The Ty1/Copia family (*Pseudoviridae*) is a widely studied group of long terminal repeat (LTR) retrotransposons present in diverse eukaryotes that share a common ancestor with retroviruses. The founding member Ty1 of *Saccharomyces* assembles into heterogeneous virus-like particles (VLPs) that are composed of the Gag and Gag-Pol proteins. We used cryo-electron tomography (cryo-ET) of Ty1 VLPs purified from a Ty1-less yeast strain expressing a functional Ty1 element to determine the organization and interactions of Gag in Ty1 VLPs. Using sub-tomogram averaging (STA), we observed unique capsomere structures and obtained EM density maps for pentagonal (7.5 Å) and hexagonal (7.9 Å) capsomeres. Through iterative rounds of alignment and classification, we demonstrate that Ty1 VLPs are composed of diverse arrangements of capsomeres, unlike true icosahedral virions. We modeled the structure of the conserved Ty1 Gag capsid C-terminal domain (CA-CTD), previously determined using X-ray crystallography into the capsomeric arrangements, and identified roles for the Dimer-1 interface present in the asymmetric unit, as well as additional interfaces involved in intra- and inter-capsomere assembly. We show that pentagonal or hexagonal capsomere assembly results from flexibility of CA-CTD – CA-CTD interactions at Interface-2. Additionally, we determined the structure of the Ty1 Gag capsid N-terminal domain (CA-NTD) using solution NMR and fitted the CA-NTD into the cryo-ET density maps. Our results indicate that the heterogeneous population of VLP size and morphology arises, at least in part, from the ability of the capsomeres to assemble in diverse organizations. (238 words)

## Introduction

Long terminal repeat (LTR) retrotransposons are a major and widely disseminated class of transposable elements (TEs) whose gene structure, life cycle, and phylogeny parallel that of retroviruses.^1,2,3,4^ However, LTR retrotransposons lack an envelope gene, and transposition is likely not infectious.^5^ Intensive studies of the Ty1/Copia (*Pseudoviridae*) and Ty3/Gypsy (*Metaviridae*) families in *Saccharomyces* and *Drosophila* have defined key steps in the process of LTR-element retrotransposition, serving as paradigms for understanding retroelement biology.

LTR retrotransposons typically contain two open reading frames; *GAG* that encodes the structural proteins comprising virus-like particles (VLPs), including capsid (CA) and nucleocapsid (NC), and *POL* that encodes enzymes required for VLP maturation (protease, PR), integration (integrase, IN), and reverse transcription (reverse transcriptase, RT). In the case of Ty1, an archetypal LTR retrotransposon present in *Saccharomyces* spp., Pol proteins are expressed via a programmed frameshifting event, resulting in the Gag-Pol (p199) precursor protein, which is incorporated into the VLP along with the Gag (p49) precursor.^6^ Ty1 Gag contains CA and NC functionality and, along with Gag-Pol, is proteolytically cleaved to form mature p45 Gag, as well as mature PR, IN, and RT proteins derived from Pol.^7,8,9,10^ Like other LTR retrotransposons, Ty1 Gag CA consists of a capsid N-terminal domain (CA-NTD) and a capsid C-terminal domain (CA-CTD). Interestingly, Ty1 *GAG* contains a self-encoded restriction factor, p22, consisting of the Gag CA-CTD arising from a sub-genomic transcript that inhibits VLP assembly through interactions with full-length Gag.^10,11,12,13,14^ Interactions between the CA-NTD and CA-CTD domains drive VLP assembly and define particle morphology. Moreover, particle assembly is a crucial step in the life cycle of LTR retrotransposons, serving as a subcellular compartment for the enzymatic activities of PR, IN, and RT to synthesize and coordinate the pre-integration complex.

Virus and virus-like particles assemble through oligomerization of an individual asymmetrical capsid subunit into icosahedral structures. Capsids form particle sub-structures called capsomeres, and equivalent interactions between capsomeres would result in a flat sheet. To overcome strict equivalence, capsomeres form quasi-equivalent interactions, giving particles the inflection points required to form icosahedral spherical structures. In retroviruses and retrotransposons 12 pentagonal capsomeres form the inflection points that are necessary for the curvature to fully close a capsid lattice containing a variable number of hexagonal capsomeres.^15^ The number of hexagonal capsomeres in a particle defines its T-number. This quasi-equivalence is apparent in high-resolution virus particle structures for several retroviruses including murine leukemia virus (MLV), human immunodeficiency virus (HIV), Rous sarcoma virus (RSV), Mason-Pfizer monkey virus (MPMV), Prototype foamy virus (PFV) and the LTR retrotransposons Ty3 and *Copia*.^16,17,18,19,20,21,22^ In the orthoretroviruses that undergo distinct structural changes associated with particle maturation^23^ the capsid lattice of immature retroviruses is not fully closed, with gaps and lattice irregularities that allow for curvature without pentameric capsomeres.^17,19,24,25^ Notably, the retrotransposons of immature PR-defective Ty3 VLPs exhibit a high degree of flexibility, with multiple T-numbers present.^20^ Structural characterization of somatic nuclear populations of *Copia* VLPs reveals homogenous T=9 icosahedral structures.^22^ By contrast, negative stain electron microscopy and low-resolution cryogenic electron microscopy (cryo-EM) studies show that full-length and C-terminally truncated Ty1 Gag assembles into VLPs with a high degree of size and morphological variation.^26,27,28^

The unique functions of LTR retrotransposon Gag and CA, including their propensity to assemble into VLPs containing element RNA and ubiquitous presence as repeated sequences in distantly related organisms, have led to a growing number of host domestication events and repurposing Gag/VLPs for cellular functions.^29,30,31^ Notably, several domesticated LTR retrotransposon Gag proteins with critical cellular roles have been characterized functionally and structurally. The neuronal gene *Arc* is a master regulator of synaptic plasticity and neuronal function and was endogenized independently in the fly and tetrapod lineages.^32,33,34,35,36^ dArc1 forms T=4 icosahedral structures with CA-NTD and CA-CTD domains forming contacts necessary for oligomerization.^14,37^ The endogenously expressed Gag gene, which encodes the paraneoplastic Ma antigen (PNMA), is associated with paraneoplastic syndrome in humans, a condition in which the immune system is activated in response to antigens presented by cancerous cells.^38^ PNMA proteins contain CA-NTD and CA-CTD domains, assembling into T=1 icosahedral structures that trigger immune responses and neuronal deficits.^39,40^ *PEG10* is a mammalian LTR retrotransposon-derived gene that encodes a Gag protein containing CA domains, binds RNA, is involved in placental formation, and assembles into particles that can modulate gene expression in neighboring cells.^41,42,43,44^

The mechanisms underlying retrotransposon VLP size and morphological heterogeneity, as well as whether VLPs display proper icosahedral symmetry, are incompletely understood. Here, we characterized the organization of transposition-competent Ty1 VLPs^5,45^ using a combination of structural approaches. Ty1 VLPs from the active Ty1H3 element were purified, and sub-tomogram averaging from cryogenic electron tomography (cryo-ET) data sets revealed several classes of capsomere organization. To better understand the oligomeric properties of Ty1 Gag, we solved the CA-NTD domain using solution nuclear magnetic resonance (NMR). We fit the CA-NTD along with the previously determined crystal structure of the CA-CTD into our cryo-ET density maps. Our findings elucidate the structural basis for Ty1 VLP heterogeneity and broaden our understanding of LTR retrotransposon VLP assembly.

## Results

### Ty1 VLPs are heterogeneous in size

To determine the Ty1 VLP structural organization, we performed cryo-ET of purified particles (See Materials and Methods and **Supplemental Table 1** for details). To ensure VLP heterogeneity was not due to Ty1 Gag sequence diversity, we purified VLPs from a Ty1-less *Saccharomyces paradoxus* strain expressing the active Ty1H3 element fused to the *GAL1* promoter on a multicopy plasmid (**Figure 1A**).^5,45,46,47^ VLPs assembled *in vivo* were purified through a series of sucrose and iodixanol gradients (**Figure 1B and C**). SDS-PAGE analysis of purified VLPs reveals prominent bands corresponding to Ty1 Gag (∼50 kDa) and additionally Killer L-A Gag (∼76 kDa) (**Figure 1D**). Ty1 Gag exists as both p45 and p49 protein products and may be glycosylated, resulting in multiple species with an approximate molecular weight of 50 kDa.^12,48^ Slices from cryo-ET tomograms of Ty1 VLPs revealed approximately spherical particles with significant heterogeneity in particle size and morphology (**Figure 1E and F**), consistent with previous studies.^27,28^ Ty1 VLPs have a rough surface with punctate electron-dense areas, and some VLPs appear to have a broken or open morphology. Killer L-A virus particles co-purified with Ty1 VLP preparations^49,50^ and were present in our VLP cryo-ET data sets as regularly sized particles with a sharper surface (**Figure 1E and G**). Killer L-A virus particle morphology is sufficiently distinct from Ty1 VLPs to easily distinguish between the two in raw micrographs, allowing us to train particle picking on only Ty1 VLP particles. Confirmation of the identity of the Killer L-A virus particles was performed by sub-tomogram averaging of the particles. The resulting map (4.4 Å; FSC @ 0.143) was consistent with a previously determined structure of L-A virus (**Supplemental Figure 1**).^49^ Distinct morphological changes are observed in retroviruses and the Ty3 LTR retrotransposon upon particle maturation.^19,20,51^ However, we did not observe any distinct morphological changes that might correspond to maturation of Ty1 Gag p49 to p45, contrary to previous observations of two distinct particle size populations of Ty1 VLPs.^28^

**Figure 1.**
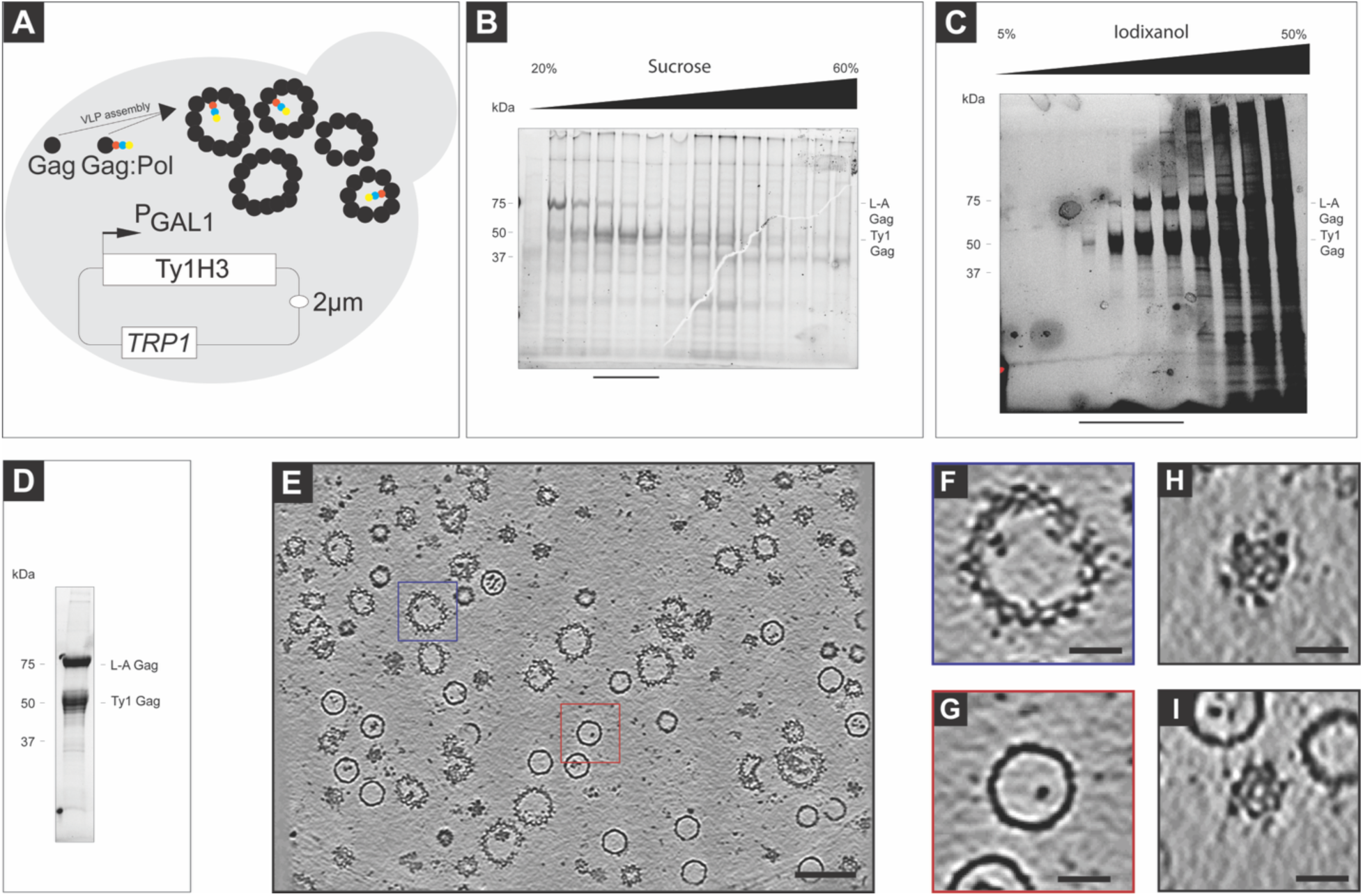
Cryo-ET of purified Ty1 VLPs. (**A**) Schematic representation of Ty1 VLP production in yeast strain DG4226. The Ty1H3 element was expressed with the GAL1 promoter present on a high-copy 2 μm *TRP1* plasmid. Ty1 Gag and Gag-Pol proteins assemble into VLPs *in vivo*. (**B**) SDS-PAGE of 20% to 60% (w/v) sucrose gradient ultracentrifugation fractions containing Ty1 VLPs. Fractions enriched for Ty1 Gag, indicated by the black bar beneath the lanes, were pooled and used for iodixanol gradient fractionation. (**C**) SDS-PAGE of 5% to 50% iodixanol gradient ultracentrifugation fractions containing Ty1 VLPs. Fractions enriched for Ty1 Gag, indicated by a black bar beneath the lanes, were pooled for buffer exchange into VLP buffer A. (**D**) SDS-PAGE of final pooled iodixanol gradient fractions following buffer exchange. (**E**) Single slice from the IsoNet processed tomogram of Ty1 VLPs purified from yeast strain DG4226. Both Ty1 VLPs (blue box, panel F) and Killer L-A virus (red box, panel G) capsids are present as distinct populations in the tomograms. (**F**) Enlarged image of a Ty1 VLP cross-section (blue box in E). (**G**) Enlarged image of a Killer L-A virus capsid cross-section (red box in A). (**H**) Enlarged image of a Ty1 VLP capsomere that appears to have six neighboring capsomeres from a different z-slice of the tomogram in A. (**I**) Enlarged image of a Ty1 VLP capsomere that appears to have five neighboring capsomers from a different z-slice of the tomogram in A. Scale bar 100 nm in E, 25 nm in F, G, H, I.

### Ty1 VLPs display varied capsomere organization

After processing the tomograms with IsoNET to denoise and correct for the missing wedge, differences in the organization of capsomeres became apparent in the tomograms. Some regions of the VLPs appeared pentameric (**Figure 1H**), while others were hexameric (**Figure 1I**). However, due to the high degree of heterogeneity in the size and morphology of Ty1 VLPs, averaging of full VLPs was not possible. Therefore, we utilized sub-tomogram averaging (STA) to determine the structure of smaller regions of the VLPs. EMAN2 was used to segment and then generate initial model points on Ty1 VLPs (see Materials and Methods). Initial rounds of STA utilized the same initial points but aligned using either a pentameric or hexameric capsomere as a single-particle initial reference. PCA classification was used to classify the particles after initial alignment. Through multiple rounds of alignment and classification (**Supplemental Table 2**), we identified numerous capsomere organizations within the VLPs (**Figure 2A-Y**). We identified a pentameric capsomere surrounded by hexameric capsomeres, an organization also seen in virions with true icosahedral symmetry (**Figure 2A-E**). Interestingly, we also resolved a structure of neighboring pentameric capsomeres surrounded by hexameric capsomeres that is incongruent with icosahedral symmetry (**Figure 2F-J**). A major structural class centered on a hexameric capsomere had two opposing pentameric subunits surrounding it, similar to the organization along the C2 axis of a T=4 icosahedral particle (**Figure 2K-O**). We also determined averages for hexameric capsomeres with one (**Figure 2P-T**) or no (**Figure 2U-Y**) neighboring pentameric capsomeres. This data demonstrates that Ty1 VLPs are comprised of both pentameric and hexameric subunits, and that these subunits assemble into a variety of arrangements relative to one another, unlike true icosahedral virions. The capsomere organizational variation observed is consistent with the heterogeneity in overall size reported here and in earlier studies of Ty1 VLP structure.^26,27,28^

**Figure 2.**
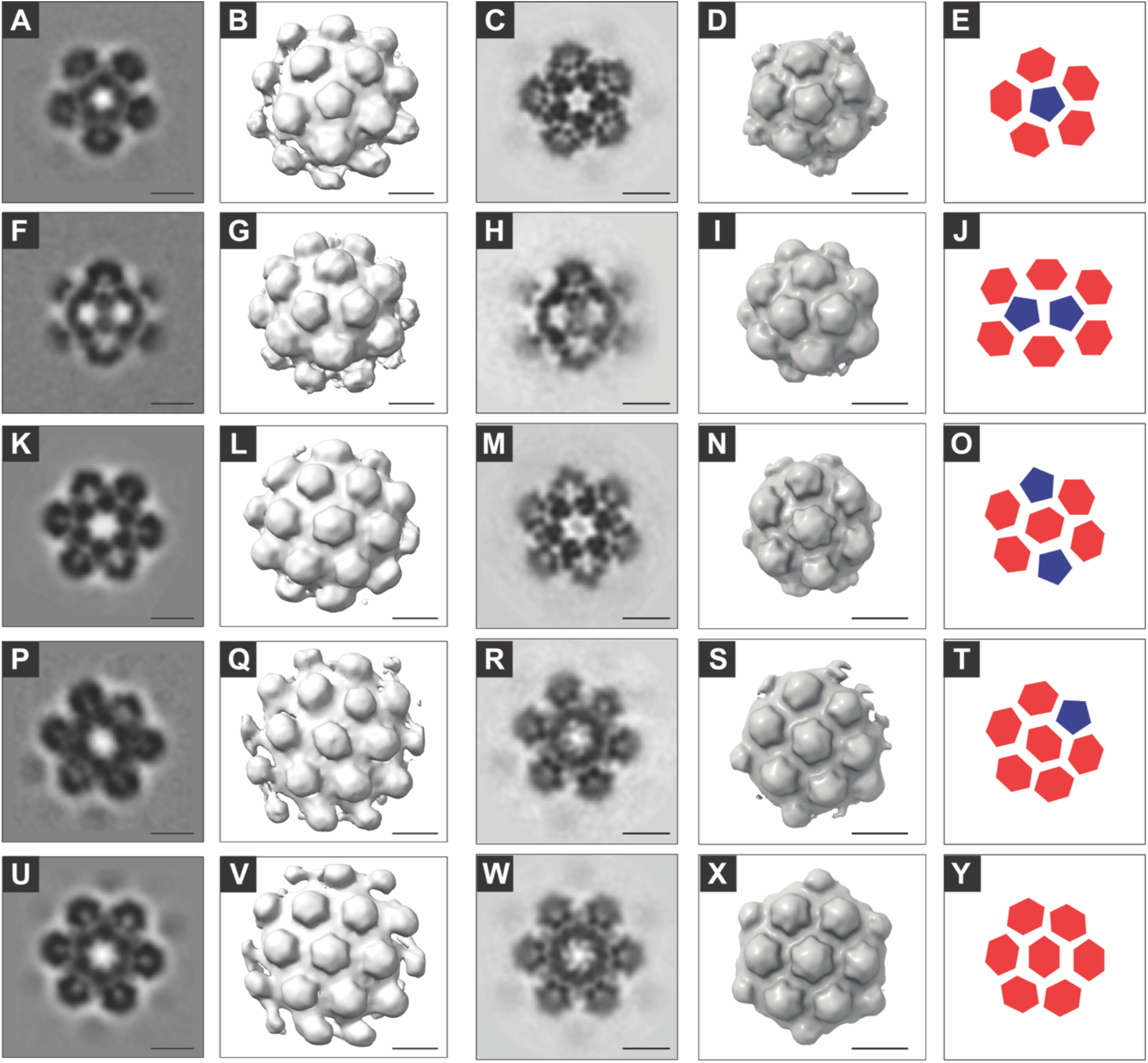
Sub-tomogram averages of Ty1 VLP capsomeres organization. (**A**) Single slice from an electron density map of a sub-tomogram average from Ty1 VLPs centered on a pentameric capsomere following multiple rounds of alignment and PCA classification. Final alignment was performed in PEET at a bin size of 4 (6.77 Å/px), using 6064 particles. (**B**) Isosurface of the density map in A. (**C**) Single slice from electron density map of a sub-tomogram average of particles in **A** aligned and reconstructed in Relion at bin4. (**D**) Isosurface of map in (C). (**E**) Cartoon depiction of capsomere arrangement based on averages in PEET and Relion. Hexameric capsomeres are shown in red, and the central pentameric capsomere is shown in blue. (**F**) As in A, showing the subtomogram average from a different set of particles containing two neighboring pentameric capsomeres surrounded by hexameric capsomeres. Final alignment includes 3582 particles. (G, H, I, J) As described for B, C, D, E, respectively, for the particles included in panel F. (K, L, M, N, O) as in A, B, C, D, E showing a hexameric capsomere with two pentameric capsomers on opposing sides. Final alignment includes 13,616 particles. (P, Q, R, S, T) As in A, B, C, D, E showing a hexameric capsomere surrounded by a single pentameric capsomere and five hexameric capsomeres. Final alignment includes 3,327 particles. (U, V, W, X, Y) As in A, B, C, D, E showing a hexameric capsomere surrounded by six hexameric capsomeres. Final alignment includes 4,724 particles. Scale bars are 10 nm.

### Ty1 Gag CA domain organization within capsomere densities

To better understand the interactions of individual copies of Ty1 Gag within pentameric and hexameric capsomeres, we determined higher-resolution structures through additional refinement. The pentameric capsomere average (**Figure 2C**) was used as an initial reference for template matching across 87 tomograms using PEET and RELION 4.0 (**Supplemental Table 2**). Through iterative rounds of refinement, we resolved a 7.52 Å pentameric capsomere structure (**Figure 3A-D, Supplemental Movie 1, Supplemental Figure 2**). The hexameric capsomere average (**Figure 2M**) was used as an initial reference for a second set of template matching across the same 87 tomograms using PEET and RELION 4.0. The final resolution of the hexameric capsomere was 7.95 Å (**Figure 3E-H, Supplemental Movie 2, Supplemental Figure 3**). In both higher resolution capsomere averages, we observed an outer ring of more ordered densities with lower ordered densities in the central turret (**Supplemental Figure 4**). Because we were unable to resolve side chains or secondary structures at this resolution, we sought to obtain more detailed information on the organization of individual copies of Ty1 Gag within capsomeres by fitting higher resolution atomic models solved by NMR and X-ray crystallography.

**Figure 3.**
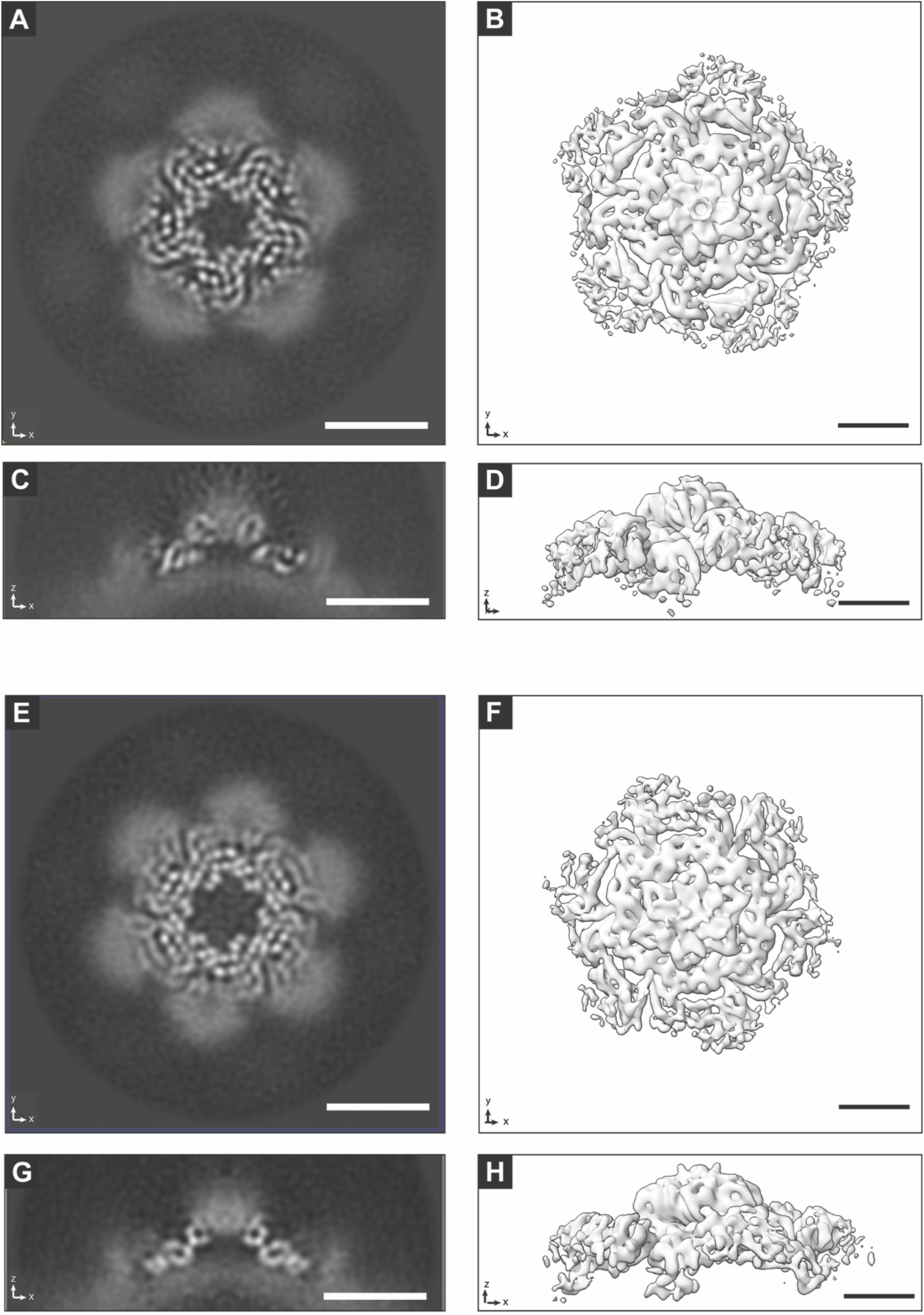
Sub-tomogram average of Ty1 VLP capsomeres. (**A**) Single X, Y slice from electron density map from sub-tomogram averaging of a pentameric capsomere from Ty1 VLPs. (**B**) Isosurface of the map in A. (**C**) Single X, Z of electron density map in A. (**D**) Isosurface in A rotated 90° around the X-axis, matching the orientation in C. (**E**) Single X, Y slice from the electron density map from sub-tomogram averaging of a hexameric capsomere from Ty1 VLPs. (**F**) Isosurface of map in E. (**G**) Single X, Z of electron density map in (E). (H) Isosurface in B rotated 90° around the X-axis, matching the orientation in G.

### Solution NMR structure of Ty1 CA-NTD

To model Ty1 CA-CTD we used the previously determined X-ray structure of p18m^14^ that has the identical sequence to Ty1 CA-CTD residues (259-355). As no model was available for placing the other regions of Ty1 Gag, we determined the solution structure of Ty1 CA-NTD, (residues T163-Q260) using NMR. The 2D ^15^N-^1^H and ^13^C-^1^H NMR spectra obtained were highly dispersed and well resolved (**Supplemental Figure 5**) and we obtained a final model calculated based on 2251 distance restraints, 72 pairs of backbone torsion angle restraints and 46 hydrogen bond distance restraints (**Supplemental Table 3**). The family of the 20 lowest-energy conformers is well defined (**Supplemental Figure 6**), with an average root-mean-square deviation (RMSD) from the mean backbone coordinates of 0.22 ± 0.05 Å and 0.59 ± 0.09 Å for all residue heavy atoms in the core 4-helix bundle. The remaining residues in loop regions, particularly the long loop that links α1-α2, do not have well-defined positions and additionally have regions undergoing µs-ms conformational exchange as judged from the NMR relaxation data (**Supplemental Figures 7 & 8**). Nevertheless, there are no restraint violations > 0.5 Å in the core structure and only small deviations from ideal geometry confirming the good quality of the final structure (**Supplemental Table 3**).

The Ty1 CA-NTD structure (**Figure 4A**), contains an N-terminal extended region and a tightly packed antiparallel four α-helix bundle. The four α-helices consist of residues D180 to S195 (α1), D214 to F227 (α2), T235 to V243 (α3) and I248 to M259 (α4), with helix α1 and α2 in an orientation that is approximately perpendicular to that of α3 and α4. The side chains from the N-terminal extended region (residues V169-L175) pack into a cleft formed between helices α1 and α2 of the bundle. A long loop is situated between α1 and α2, and two short loops connect α2-α3 and α3-α4. This twisted arrangement of the four secondary structure elements forms a hydrophobic core that is stabilized by a variety of hydrophobic interactions involving the aliphatic and aromatic sidechains that are located on the inner faces of the helices. These tertiary interactions that involve aromatic sidechains give rise to several readily identifiable ^1^H chemical shift outliers due to ring current effects. Most notably, the backbone amide proton (6.00 ppm), Hα (2.96 ppm) and Hγ2 methyl group of V237 (1.30 ppm) display large upfield shifts (**Supplemental Figure 5**) because of the proximity to the aromatic sidechains of F181, F224 and W236. Upfield ^1^H chemical shifts were also identified in ^13^C-^1^H correlation spectrum (**Supplemental Figure 5B**) for the ^1^Hβ methylene group (-0.28 ppm) and ^1^Hψ (0.40 ppm) of L175, which is positioned in between the aromatic side chains of F181 and W184, for the ^1^Hψ2 methyl group of I226 (0.02 ppm), which is oriented towards F227 and for methyl group of A228 (-0.32 ppm) which faces the aromatic rings of F181, W184 and F224.

**Figure 4.**
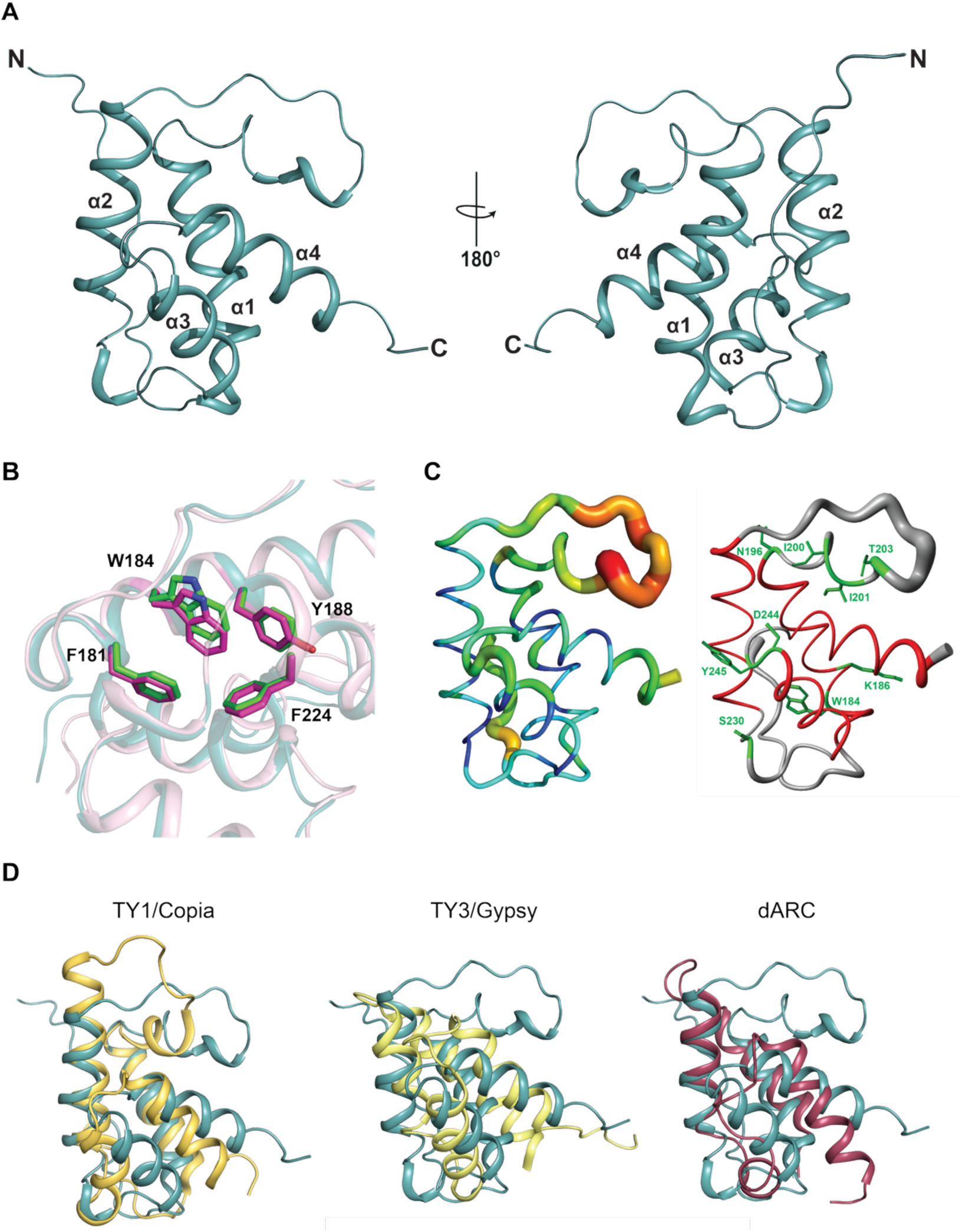
Ty1 CA-NTD solution NMR structure. **(A)** Lowest energy solution NMR structure of Ty1 CA-NTD. The backbone is shown in light teal cartoon representation in two views rotated by 180°, α-helices are labelled sequentially from the N- to the C-terminus. (**B**) Close-up view of the superposition W184 and surrounding residues F181, Y188 and F224 in the major (green) and minor (magenta) conformations of TY1 CA-NTD. The protein backbones are shown transparent cartoon, major conformer in teal and minor in magenta. The W184 rotamer of the major conformation has *χ*1=176° and the minor conformer has *χ*1 = 167°. (**C**) 3D representation of Ty1 CA-NTD dynamics. (Left) protein putty representation, the tube represents the protein backbone with diameter at any position relating to the reciprocal value of the Het-NOE (950 MHz) for each residue. A thicker tube and coloring blue to red indicates increased motional amplitude and fast dynamics. For clarity the first seven N-terminal and two C-terminal residues that have high dynamics have been excluded. For proline residues where data is not available, the average values from the neighboring residues has been applied. (Right) protein putty representation of the spatial RMSD of 20 Ty1 CA-NTD NMR structures. The tube thickness is directly proportional to the RMSD of the ensemble. The protein backbone is colored by secondary structure, with helices in red, coil and turn in grey. Residues affected by dynamics on the µs-ms timescale as determined by ^15^N R_2_^eff^ relaxation dispersion experiments are shown in sticks, labeled and colored in green. (**D)** Structural alignment of Ty1 CA-NTD with related Gag CA-NTD domains. Pairwise PDBeFold 3D Cα structural alignment of Ty1 CA-NTD with (left) *D. melanogaster* Ty1-*Copia* CA-NTD (PDB ID: 8VVW), (center) Ty3/Gypsy CA-NTD (PDB ID 6R22) and (right) Drosophila neuronal dARC1 (pdb-code 6TAP). In each panel, the cartoon of the Ty1 CA-NTD is shown in light teal.

### Ty1 CA-NTD backbone dynamics

During the analysis of NMR spectra, we observed a set of minor peaks (∼20), each associated with a major amide resonance in the ^1^H-^15^N 2D correlation spectrum (**Supplemental Figure 5**). Backbone resonance assignment experiments enabled us to partially establish the sequential connectivities for these minor peaks and revealed them to result from duplicate assignments for a small set of resonances belonging to regions of the polypeptide chain showing the structural heterogeneity. This peak-doubling suggested there were inherent conformational dynamics within the domain arising from the presence of two species in equilibrium on a slow-exchange timescale. Quantification of peak volumes excluding significantly overlapped and severely broadened resonances, gave an average volume ratio between the minor and major species of 0.7:0.3 indicating that 30% of the protein adopts the minor conformational state. Inspection of the residues that exhibit peak-doubling revealed that many including M174, L175, N183, W184, V185, K190, F191, L192, and A228, all locate in proximity with the indole-ring sidechain of W184. In the structure, the W184 sidechain is positioned in the hydrophobic core surrounded by the aromatic rings of F181 and Y188. This prevents continuous reorientation around the Cβ-Cγ bond and sets the W184 sidechain *χ*1 dihedral angle to an average value of 176°. However, in the minor conformation a rearranged packing and a movement of W184 ring about its Cα-Cβ bond (**Figure 4B**) now results in a *χ*1 dihedral angle of 167°. Therefore, this repositioning between conformations both provides a molecular interpretation of the resonance peak-doubling and the presence of slow dynamical processes within the Ty1 CA-NTD.

### ps-ns timescale dynamics

To determine whether Ty1 CA-NTD backbone dynamics contributed to the observed major and minor conformations, we performed backbone ^15^N R_1_ and R_2_ relaxation rate and the steady state heteronuclear ^1^H-^15^N NOE, (Het-NOE) measurements at 600 and 950 MHz on uniformly ^15^N-^13^C-labeled protein. These measurements are sensitive to internal mobility and provide information on the rapid (ps-ns) motions of individual residues, as well as inform on the presence of slower (µs-ms) protein motions.^52^ Relaxation parameters were obtained for non-overlapping resonances of 82 non-prolyl residues of Ty1 CA-NTD and are plotted for each residue in **Supplemental Figure 7**. As is commonly observed for folded proteins, the Ty1 CA-NTD R_1_ and R_2_ relaxation rates away from the protein termini, are uniform with little variation across the secondary structure elements, a pattern consistent with a single overall tumbling rate for the protein. Moreover, residues within α-helices demonstrated high Het-NOE values, average of 0.8 (**Supplemental Figure 7**), indicative of the limited motion of N-H bond vectors characteristic of folded proteins.^53,54^ The N- and C-terminal residues display much larger R_1_ and much smaller R_2_, identifying regions of the protein with increased local flexibility. However, in contrast to stable secondary structure regions, R_1_ for residues within the long α1-α2 interconnecting loop gradually increases (T203-K207) then decreases after P208 (V209-I212). R_2_ shows the opposite effect, sharply decreasing from T203 to K207 then increasing from V209-I212, behavior indicative of the fast motions of highly flexible residues within this loop. In addition, the Het-NOE values are reduced ∼0.6, which is smaller than that observed for well-ordered proteins^55^ indicating the presence of large amplitude NH bond vector motions. Taken together, these data suggest that in addition to the overall molecular tumbling, the loop region comprising residues T203-I212 experiences fast local motion on the ps-ns timescale that is consistent with a dynamic environment and their uncertain positioning within the NMR ensemble (**Figure 4C & Supplemental Figure 6**). Structural similarity searches of Ty1 CA-NTD performed using the PDBeFold search engine (https://www.ebi.ac.uk/msd-srv/ssm/) identified the CA-NTD from the *D. melanogaster* retrotransposon *Ty1-Copia* CA-NTD as the top hit based on Z score (**Supplemental Table 4**) followed by much weaker hits with the CA-NTD from retrotransposon *Ty3/Gypsy* and that of the CA-NTD from the exapted Gag protein of *D. melanogaster* neuronal protein dARC1. Pairwise 3D Cα structural alignment of Ty1 CA-NTD (**Figure 4D**) show that whilst the CA-NTDs maintain the same four-helix bundle topology, there is significant variation in helix crossing angles and packing within each domain. In addition, the α1-α2 long loop of Ty1 CA-NTD is much shorter in the *Ty1-Copia* CA-NTD and largely absent from the CA-NTD of Ty3 and dARC, suggesting the fast dynamical behavior we observe is likely reduced in other retrotransposon CA-NTDs.

### ^15^N NMR R2 Relaxation Dispersion µs-ms timescale dynamics

Within the relaxation data it was also apparent that residues I200 and Y245 displayed elevated R_2_ values (**Supplemental Figure 7**) indicative of conformational exchange on the µs-ms timescale. Therefore, to investigate conformational dynamics of Ty1 CA-NTD on this timescale, we applied ^15^N TROSY Carr-Purcell-Meinboom-Gill (CPMG) relaxation dispersion spectroscopy at two static field strengths of 14.1 T and 22.3 T. These experiments probe exchange dynamics between species with distinct chemical shifts on a timescale ranging from 100 µs to 10 ms.^52^ Several Ty1 CA-NTD residues did show substantial exchange contributions (>3 s^-1^) to the ^15^N transverse relaxation rates (R_ex_) (**Figure 4C & Supplemental Figure 8**). In the α1-α2 long loop residues N196, I200, I201, and T203, display dispersion curves that are characteristic of the slow exchange limit. Therefore, these dispersion data were statistically best fit with the Carver-Richards equation^56^ with global P_b_ and k_ex_ values of 0.018 ± 0.006 and 1596 ± 177 s^-1^. Residues S230, D244, and Y245 located in the α2-α3 and α3-α4 loops respectively, also display slow-exchange that was best fit with global P_b_ and *k_ex_* values of 0.010 ± 0.002 and 1162 ± 178 s^-1^ whilst residues W184 and K186 were best fit with P_b_ and kex values of 0.017± 0.0062 and 1676± 365 s^-1^. Taken together, these data demonstrate that within Ty1 CA-NTD the N-terminal portion of the α1-α2 loop and residues surrounding the W184 are undergoing slow exchange processes. These motions likely contribute to conformational switching that results in the major and minor conformations we have observed. Also, the structure shows a degree of plasticity rather than adopt one static configuration.

### Model docking into cEM maps

We rigidly docked the CA-NTD NMR structure (**Figure 4**) and the CA-CTD X-ray structure^14^ into our cryo-ET maps to determine the general orientation and organization of Ty1 Gag within the VLPs. Ty1 Gag CA-CTD residues 259-351 comprises five ordered α-helices packed into an antiparallel bundle, and this structure was solved to a resolution of 2.8 Å by X-ray crystallography.^14^ The crystallographic asymmetric unit comprised three copies of Ty1 Gag CA-CTD with two dimer interfaces (Dimer-1 and Dimer-2) defined by buried hydrophobic interactions. A biological assembly comprised of a single dimer across the Dimer-1 hydrophobic interface was assigned based on evidence from light scattering and equilibrium centrifugation (PDB ID 7NLH). It was also proposed that Dimer-2 may serve as a trimer interface in the context of a VLP.^14^ Here, the pentameric capsomere cryo-ET map suggested that five copies of the proposed biological assembly (ten individual copies of Ty1 Gag CA-CTD) fit into a ring surrounding the central turret (**Figure 5A and B**). The positioning of this structure indicated that the central turret contained residues N-terminal to the CA-CTD. Although this region is less well-resolved in the map (**Supplemental Figure 4**), we were also able to fit five copies of the Ty1 CA-NTD (**Figure 2**) into the density. The region of Ty1 Gag N-terminal of the CA-NTD, residues 1-162, is predicted to be disordered, containing a prion-like domain^57^ and was not resolved in our cryo-ET maps. Consistent with our findings of the CA-NTD occupying the central turret of the capsomere, immunological studies suggest that epitopes of the Gag CA-NTD are present on the surface of VLPs (**Figure 5B and D**).^58^

**Figure 5.**
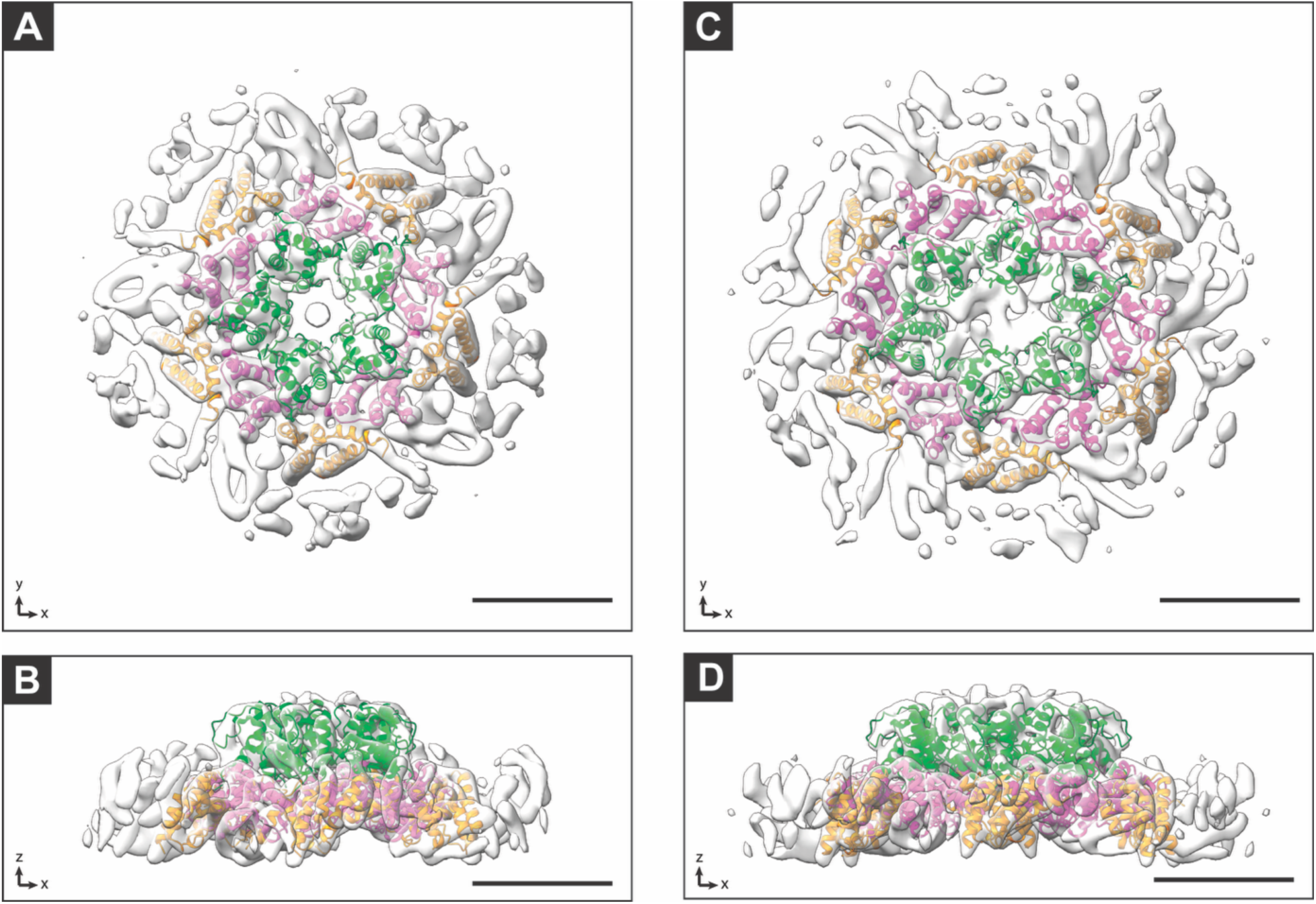
Gag CA fitting into the sub-tomogram average of Ty1 VLP capsomeres. (**A**) Pentamer isosurface shown with atomic models from Ty1 Gag CA-CTD (PDB 7NLH) (aa M259-N355) fitted into the density. Chains A (magenta) and B (tangerine) of CA-CTD were fitted together using the Dimer-1 configuration for a total of ten copies of Ty1-CA-CTD with an additional five, copies of the Ty1 Gag CA-NTD, shown in green also placed into the isosurface. (**B**) Isosurface and atomic models in A rotated 90° around the X-axis. (**C**) Hexamer isosurface with twelve copies of Ty1 Gag-CTD and six CA-NTD fitted in as in **A**. (**D**) Isosurface and atomic models in C are rotated 90° around the X-axis.

As in the pentameric capsomere, we were also able to rigidly dock the CA-CTD and CA-NTD models into our hexameric capsomere cryo-ET map (**Figure 5C and D**). Here, the proposed biological assembly, Dimer-1, fits into this map in the same general location as in our pentameric capsomere map. These observations suggest that the Dimer-1 interface is conserved in both pentameric and hexameric subunits and that the VLP heterogeneity arises from the flexibility of other interactions.

### Interactions of the Ty1 CA-NTD in capsomeres

The Ty1 CA-NTD fits into both the pentameric and hexameric capsomeres in distinct, but similar orientations (**Figure 6**). Fitting of the CA-NTD structure into the maps reveals contacts between CA-NTD copies and slight differences in the density between the pentameric and hexameric capsomeres. Though the best fit of the CA-NTD for both maps is roughly the same position, the Q-score, a measure of atomic model fit to a density map^59^, is 0.14 for the hexameric map and 0.11 for the pentameric map, indicating a better fit for the hexameric map (**Supplemental Figure 9**). Notably, the NMR spectra revealed the presence of a minor conformation in the CA-NTD and ^15^N relaxation data demonstrated both fast ns-ps and slow µs-ms dynamics, indicating significant conformational plasticity in the structure. These results suggest that the CA-NTD can adopt different conformations, with the average NMR ensemble structure more closely matching the conformation in the hexameric capsomere than that adopted in the pentameric capsomere. The local resolution of the CA-NTD is worse than that of the CA-CTD in both maps (**Supplemental Figure 9**); higher-resolution maps of this region would be required to determine differences in CA-NTD conformation within the capsid more accurately. Our NMR model was determined using the truncated CA-NTD expressed in *E. coli,* compared to our cryo-ET maps of whole VLPs isolated from a native host. The positioning of the model within our map suggests the remaining N-terminal residues (1-261) are all positioned together towards the center of the turret. We could not discern from our map if they fold back into the interior or our exterior to the VLP. The fact that they are not resolved suggests they are likely flexible or exist in multiple conformations.

**Figure 6.**
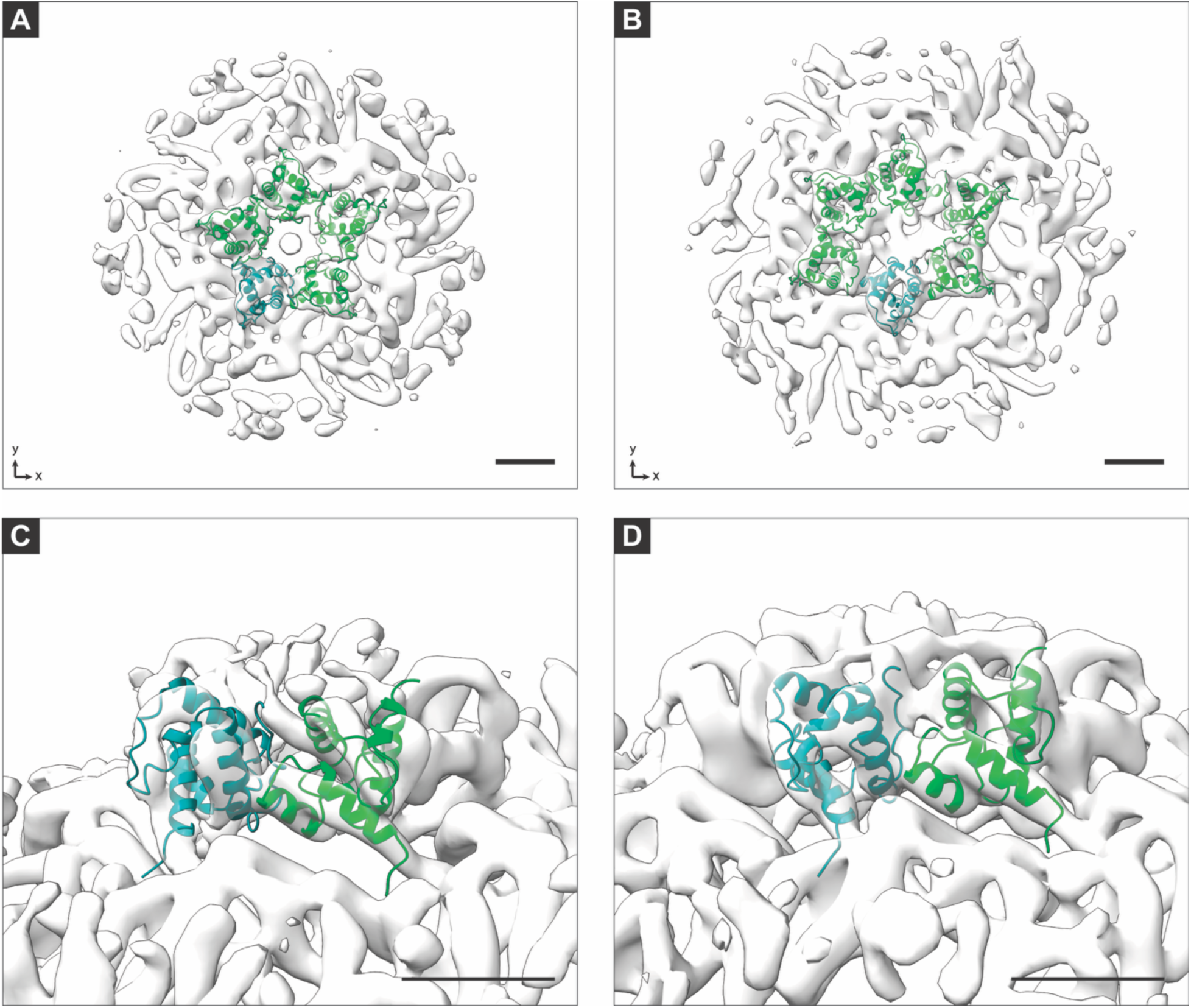
Fitting of CA-NTD into sub-tomogram averages of Ty1 VLP capsomeres. (**A**) Sub-tomogram average of a pentameric capsomere with 5 copies of CA-NTD, shown in green and teal fitted into the central turret. (**B**) As in A, the sub-tomogram average of the hexameric capsomere, with 6 copies of the NMR structure of the CA-NTD, is fit into the central turret in green and teal. (**C**) A closer view of two neighboring copies of the CA-NTD fitting into a pentamer. (**D**) As in **C**, with the hexameric capsomere average of the pentameric capsomere. Scale bars are 10 nm.

Several amino acid substitutions in the CA-NTD region are defective for retrotransposition or VLP assembly.^11^ We mapped the location of these mutations on individual copies of the CA-NTD fitted into our cryo-ET density maps (**Supplemental Figure 10**). The W184A mutation does not affect Gag protein levels but results in a loss of detectable transposition and causes altered VLP sedimentation patterns in sucrose gradients. In addition, this tryptophan residue is highly conserved in *Pseudoviridae Gag* genes.^60^ Together with our observations of the W184 sidechain engaging in hydrophobic interactions and conformational switching (**Figure 4B)**, the results reveal a key role for residue W184 in VLP assembly. Relative to a wild-type full-length Ty1H3 element, P173L and K250E substitutions result in lower fitness of an individual element but an increased resistance to p22-mediated copy number control.^11^ N283D, K186Q, and I201T substitutions result in similar fitness of an individual element and increased resistance to p22 copy number control. These mutations could be altering key CA-NTD interactions that direct VLP assembly and influence the Dimer-1 CA-CTD interface where p22 binds to Gag and restricts retrotransposition.^14^ The double mutant I248M/N249R causes an increase in the VLP diameter to 200-300 nm, another example of rearrangements of CA-NTD interactions resulting in changes in VLP assembly.^61^

### The Ty1 Gag CA-CTD forms multiple interactions in capsomeres

Fitting the Ty1 Gag CA-CTD into our sub-tomogram average of the pentameric and hexameric capsomere cryo-ET maps revealed the position of the CA–CTD – CA–CTD Dimer-1 interface as well as another three-fold interface. (**Figure 7A and B**). The Dimer-1 interface is characterized by packing of residues on α-helices 1 and 3 against residues on the neighboring α-helices 1 and 3, forming a predominantly hydrophobic buried surface. The Dimer-1 interface appears to form very similar interactions in both the pentameric and hexameric capsomeres (**Figures 7C and D**). Mutations at Gag residue 273 in α-helix 1 result in an escape of restriction from p22.^11,14^

**Figure 7.**
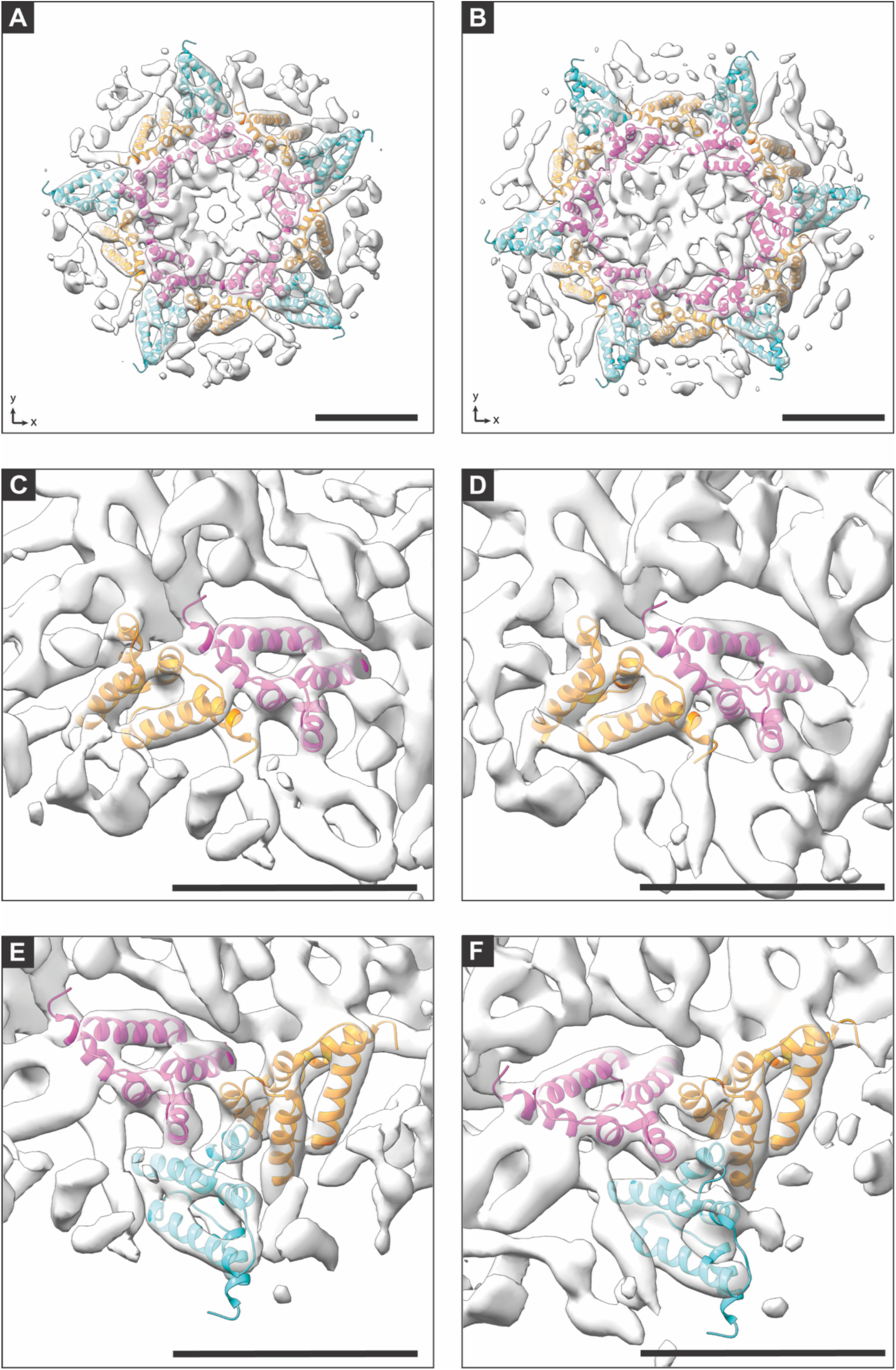
Fitting of CA-CTD into sub-tomogram averages of Ty1 VLP capsomeres. (A) The isosurface of the Ty1 pentameric capsomere, with the p18m model (PDB ID 7NLH), fitted into the map density. Fifteen copies of Ty1 CA-CTD (PDB ID 7NLH) are shown, that include dimer interfaces A (magenta) and B (tangerine) chains surrounding the central capsomere, and an additional copy of chain A (cyan), which fits into the neighboring density. (**B**) Same as in A for the hexameric capsomere. (**C**) A closer view of panel A showing model fitting of the p18m (PDB ID 7NLH) Dimer-1 interface. The model was docked with the rigid docking of chains A & B together. (**D**) As in **C** for the hexameric capsomere. (**E**) A closer view showing the interaction of three copies of p18m (PDB ID 7NLH) across Interface-2. (**F**) As in E for the hexameric capsomere. The scale, position, and rotation of the maps in Figure F were manually aligned so that the tangerine subunits of E and F are in equivalent positions. Scale bars are 5 nm.

Unlike the Dimer-1 protomer, the two monomers comprising Dimer-2 in the CA-CTD crystal structure could not be rigidly fit together in either map. However, fitting of the Dimer-1 protomer revealed an additional threefold interaction, herein referred to as Interface-2.^14^ Interface-2 is defined by three copies of the CA-CTD interacting through α-helices 4 and 5 contacts and consists of intra- and inter-capsomere contacts (**Figure 7E and F**). In contrast to Dimer-1, the orientation of the CA-CTD monomers in Interface-2 was different in the pentameric and hexameric maps (**Figure 7E and F**). Flexibility in this interaction interface thus contributes to or allows for the heterogeneous organization of Ty1 VLPs. There are likely additional conformations of Interface-2 that are not captured in our two high-resolution structures but are present in the other organizations we observed (**Figure 2**) and in additional organizations that may be present within the entire capsids.

Previous mutational analyses of residues mapping to the CA-CTD have highlighted its role in VLP assembly and transposition.^11,61^ We mapped these mutations to their locations in our high-resolution cryo-ET maps (**Supplemental Figure 11**). In a C-terminally truncated Gag comprised of residues 1-381, the I343K mutation in α-helix 5 results in VLPs with a five to six times larger diameter than wild type, and the same mutation in full-length elements results in a loss of particle assembly and transposition. I343 lies within the core of the CA-CTD, and the introduction of the longer lysine charged sidechain could cause rearrangement of key CA-CTD interactions or the introduction of interactions that are absent in wild-type VLPs. The F323S/D mutations in α-helix 4 present at Interface-2 in full-length element result in several hundred- and several thousand-fold reductions in transposition frequency, respectively.^14,62^ These mutations suggest that F323 participates in key hydrophobic interactions between CA-CTD monomers at Interface-2.

## Discussion

Our cryo-ET analysis of Ty1 VLPs isolated from a natural *Saccharomyces* host revealed multiple capsomere organizations (**Figure 2**) showing that in contrast to other viral and LTR-retrotransposon-derived VLPs, Ty1 particles display an exceptionally high degree of size and morphological heterogeneity. To better understand the molecular basis for this heterogeneity, we determined the structure of the Ty1 Gag CA-NTD domain using solution NMR (**Figure 4**). We fitted this atomic model, along with the CA-CTD atomic model, previously determined by X-ray crystallography^14^ into our cryo-ET density maps of Ty1 VLPs (**Figure 5-7**). Combining multiple structural techniques in this manner allows for a detailed study of flexible macromolecular assemblies that are not accessible by any single approach.

Viral and virus-like particles with true icosahedral symmetry are typically composed of pentagonal and hexagonal capsomeres, with the pentagonal capsomeres at the 5-fold symmetry axes and the number of hexagonal capsomers spanning these points defining the T-number and position of the 3- and 2-fold symmetry axes according to “quasi-equivalence” rules.^15^ Four out of the five classes of Ty1 capsomere organization identified in the present study can be mapped to capsomere orientations in particles with predicted icosahedral symmetry. Upon mapping of these capsomere organizations to hypothetical particles with true icosahedral symmetry, we find class 1 (**Figure 1A-E**) particles with T≥3, class 2 (**Figure 2F-J**) cannot assemble into icosahedral particles, class 3 (**Figure 2K-O**) particles with T=4, class 4 (**Figure 2P-T**) particles with T≥7, and class 5 (**Figure 2U-Y**) particles with T≥9. Together, our findings imply the presence of at least T=4 and T=9 organization of capsomeres but do not distinguish between the possibility of inter-VLP and intra-VLP heterogeneity.

However, several results suggest that intra-VLP heterogeneity is present in Ty1 VLPs. If heterogeneity were wholly between VLPs in populations of true T=4 and T=9 VLPs, the distribution of VLP sizes should be discrete rather than the continuum we observe. Other virus particles with multiple T-numbers, such as Hepatitis B, assemble *in vivo* into T=3 and T=4 particles of discrete sizes based on a conformational capsid switch.^63,64^ Previous studies of Ty1 VLP structure have also reported a continuous distribution in size, consistent with our observations.^26,27,28^ These early cryo-EM analyses suggested that Ty1 VLPs did exhibit a true icosahedral organization, but based these reconstructions on models that impose symmetry, which may introduce artifacts. In addition, if entirely inter-VLP heterogeneity were present in our data sets, we would expect larger regions of VLPs to average together, which we did not observe. In the *Saccharomyces Metaviridae* retrotransposon Ty3, intra-VLP heterogeneity is present in 13% of VLPs analyzed by cryo-ET.^20^ Our results support similar intra-VLP heterogeneity of Ty1 VLPs and further characterize this heterogeneity by examining several capsomere classes. Therefore, further study is required to understand the functional significance of Ty1-VLP heterogeneity. As we characterized a variety of VLPs isolated from yeast, transpositionally competent VLPs may display specific capsomere organizations, and transpositionally incompetent VLPs may contain other “dead-end” capsomere organizations. Notably, although Ty1 VLPs protect Ty1 RNA from benzonase (∼30 kDA) but not RNase A (∼14.7 kDA) nuclease treatments^26^, our capsomere density maps do not support regular gaps in VLP structure that would allow even entry of RNase A access to packaged Ty1 RNA. However, VLPs containing intra-capsomere heterogeneity may have irregularly distributed gaps that are large enough for the entry of RNase A.

Like other viral particles, the interactions between the Ty1 Gag CA-NTD and CA-CTD domains drive VLP assembly. We observe flexibility in both CA-NTD and CA-CTD interactions. This flexibility accommodates both pentameric and hexameric capsomeres as well as the variation detected in capsomere organization (**Figures 2, 6, and 7**). Mutating residues in the CA-NTD and CA-CTD that result in VLP assembly defects and resistance to p22 restriction also support our capsomere cryo-ET maps and indicate how perturbations to the network of intra- and inter-capsomere interactions can result in VLP morphological changes (**Supplemental Figures 10 and 11**). W184 is conserved across members of the *Pseudoviridae* and previously shown to ablate transposition and alter VLP assembly when mutated to an alanine residue.^11,60^ Our NMR µs-ms timescale dynamics results reveal a W184 adopts multiple conformational states (**Figure 4 and Supplemental Figure 8)**. Together, these results suggest that W184 conformational state switching plays a key role in VLP assembly, possibly through a mechanism mediating pentameric vs hexameric capsomere formation. Furthermore, the CA-NTD monomer solution NMR structure shows significant dynamics and structural plasticity and is not entirely congruent with our cryo-ET density maps, suggesting there are different conformations when compared with the CA-NTD in the context of full-length Gag in a VLP (**Figures 4 and 6**). Nevertheless, overall, the CA-NTD atomic model aligns well with both our high-resolution pentamer and hexamer capsomere maps. Moreover, the C-terminus of the CA-NTD and the N-terminus of the CA-CTD are adjacent in our model, as determined by fitting to the cryo-ET density maps, supporting our interpretation of individual Gag copy locations.

In the CA-CTD crystal structure, the asymmetric unit consists of three monomers interacting through two dimer interfaces defined by predominantly hydrophobic surfaces: Dimer-1, consisting of α-helices 1 and 3, and Dimer-2, consisting of α-helices 4 and 5.^14^ In our cEM maps we observed the Dimer-1 interface stabilizing the individual subunit of capsomeres containing two copies of Ty1 Gag. This interface is the target of the N-terminally truncated Ty1 self-encoded p22 restriction factor, which is composed of the CA-CTD domain. Through occupation of the Dimer-1 interface in the assembling VLP, p22 blocks the necessary contact points by making dead-end interactions to prevent or poison VLP assembly.^11,12,14^ p22 acts during early VLP assembly, preventing Gag oligomerization without disrupting the steady-state levels of the monomer.^65^ Furthermore, three key residues at positions 266, 270, and 312 at the Dimer-1 interface determine p22 restriction specificity within the Ty1 family.^66^ Thus, orthogonal analyses of the Ty1 restriction factor support our findings of the position of the Dimer-1 interface within an assembled VLP (**Figure 7**). However, our results do suggest that the Ty1 Gag CA-CTD interacts at the 3-fold Interface-2 in a VLP, rather than the Dimer-2 present in the crystal structure, and again, highlights the flexible nature of Gag’s capsid interactions.^14^

Interface-2 is present in both the pentagonal and hexagonal capsomeres and contributes to the ability of Ty1 Gag to form both types of oligomeric structures. Together, our Ty1 VLP cryo-ET analyses further our mechanistic understanding of Ty1 VLP assembly and how the p22 restriction factor disrupts this process. Further work is required to elucidate how p22 traps VLP assembly intermediates into non-functional assembly pathways.

The choice of purification method and host is essential for interpreting the biological role of VLP structures. We chose to isolate Ty1 VLPs from their natural host, *Saccharomyces,* to avoid adsorption-based purification methods or purifying recombinant, affinity-tagged Gag or assembled VLPs from a heterologous host, such as *Escherichia coli.* Alternative Ty1 VLP isolation methods may have resulted in more homogeneous VLPs that are amenable to single-particle analysis. Still, these structures could be biased towards specific sub-populations and not fully representative. Therefore, our study provides an unbiased approach to determine VLP structure from other LTR-retrotransposons and endogenous retroviruses.

Several LTR-retrotransposon *Gag* genes have been domesticated in distantly related species for novel functions, and deciphering their mechanisms requires a structural understanding of their VLPs.^67,68^ The neuronal gene *Arc* plays an important role in synaptic plasticity and was independently domesticated in the fly and tetrapod lineages from the *Metaviridae* LTR-retrotransposon *Gag*. Arc proteins form VLPs that are important for function.^35,36,69^ The fly LTR-retrotransposon *Copia*, a member of the *Pseudoviridae* family along with Ty1, acts in an antagonistic manner to *dArc1*.^70^ Compelling genetic evidence supports this antagonism, but the structural basis of the antagonism remains incompletely understood. This could be due in part to conclusions drawn from comparisons with *Copia* VLPs purified from the heterologous host *Escherichia coli,* which do not fully represent *in vivo* assembled VLPs.

The flexible nature of Ty1 VLPs, particularly their significant variability in particle size, appears to be a unique feature of LTR retrotransposons. Retroviruses do not exhibit this same level of size variability, although they do show significant morphological diversity in the capsid across different genera. Crucially, in retroviruses, this diversity is primarily associated with specific, required conformational changes that occur during particle maturation.^16,17,18,19,24,25^ These maturation-specific morphological changes are distinct from the inherent size flexibility observed with Ty1 VLPs. The flexibility of Ty1 VLP assembly observed here is consistent with the hypothesis that differences in selective pressures of LTR retrotransposons and retroviruses explain the lack of distinct mature and immature particle steps.^20^ LTR retrotransposons reside entirely within the cell with their genomic RNA or cDNA continually at risk of degradation by host factors. The flexibility of Ty1 VLP assembly may well be required to rapidly encase and protect the genome. In retroviruses such as HIV, Gag drives viral egress from producer cells whilst packaging the genome within an irregular immature particle. Maturation then occurs within the enveloped virion away from the cell preventing the opportunity for host factor degradation^71^ and producing regular cores that may be required for early events in the infection of target cells.

## Materials and Methods

### Yeast strains and plasmids

Standard yeast genetic and microbiological techniques were used in this work.^72^ Yeast *Saccharomyces paradoxus* DG3582 *(MATα gal3 his3-Δ200hisG trp1 ura3 Ty-less*) was derived from the sequenced strain DG1768.^47^ Plasmid pBDG538, a multicopy TRP1 shuttle plasmid containing a full-length Ty1H3 element driven by the *GAL1* promoter, was introduced to DG3582 to generate DG4226.

### VLP purification

VLPs were purified as previously described^12^, with the addition of an iodixanol gradient and iodixanol removal by spin concentration steps. Briefly, strains DG4226 and DG4493 were grown overnight at 30 °C in 40 mL SC-Trp + 2% raffinose, diluted 1/25 into 1 L SC-Trp + 2% galactose and grown for 48 hours at 22 °C. 6 L of cells grown in SC-Trp + 2% galactose were harvested for each VLP purification. Cells were harvested by centrifugation at 4 °C, 6000 x *g* for 25 minutes. The cell pellets were resuspended in 30 mL of VLP Buffer A (15 mM KCl, 10 mM HEPES-KOH, pH 7.6, 5 mM EDTA) per 1 L of culture and cells collected by centrifugation at 4 °C at 6000 × *g* for 10 minutes. Pellets were resuspended in 3.2 mL VLP Buffer A plus protease inhibitor cocktail (Aprotinin 125 μg/mL, Leupeptin 125 μg/mL, Pepstatin 125 μg/mL, 1.6 mg/mL phenylmethylsulphonyl fluoride, and 93.75 mM Dithiothreitol) per 1 L of harvested culture. Acid-washed glass beads were added, and cells were lysed by vortexing for 16 cycles of 1 minute with 1-minute rest intervals on ice. The lysate was clarified at 12096 *x g* for 10 minutes and loaded onto a step gradient of 20%, 30%, 45%, and 70% (w/v) sucrose in VLP Buffer A. The gradient was centrifuged at 112398 *x g* for 3 hours in a SW28 rotor. 4 mL of the layer between 30% and 45% steps was collected, diluted to 10% sucrose in VLP Buffer A, and centrifuged at 277071.9 x *g* for 45 minutes in a Ti-70.1 rotor. The pellet was resuspended in 400 μL VLP Buffer A and centrifuged through a 20% to 60% continuous sucrose gradient in VLP Buffer A at 112398.1 *x g* for 3 hours in a SW28 rotor. The gradient was fractionated using an ISCO Foxy Jr. fraction collector (Lincoln, NE), and fractions were analyzed by SDS-PAGE on a 10% TGX Stain-Free gel (Bio-Rad Cat # 4568036). Peak fractions were chosen based on the intensity of the ∼ 50 kDa band corresponding to Ty1 Gag, pooled, diluted in VLP Buffer A, and centrifuged at 277071.9 x *g* for 45 minutes in a Ti-70.1 rotor. The pellet was resuspended in 200 μL of VLP Buffer A and centrifuged through 5% to 50% iodixanol gradient in 1 mM EDTA, 20 mM Tris-HCl, pH 7.4, at 213626.2 x *g* for 3 hours in a TLS55 rotor. The iodixanol gradient was fractionated manually by puncturing the tube with a 20-G needle. Fractions were analyzed by SDS-PAGE on a 10% TGX Stain-Free gel (Bio-Rad Cat # 4568036). Peak fractions were chosen based on the intensity of the 50 kDa band corresponding to Ty1 Gag, which was then pooled and exhaustively buffer-exchanged 1:4 into VLP Buffer A 11 times. The final sample was analyzed by SDS-PAGE on a 10% TGX Stain-Free gel (Bio-Rad Cat # 4568036) and used for cryo-EM and ET data collection.

### Grid preparation

Purified Ty1 VLPs were applied to glow-discharged, Quantifoil R1.2/1.3 200 mesh copper grids with 2 nm continuous carbon support. Grids were blotted for 2-4 seconds and plunge frozen into liquid ethane using a Vitrobot Mark IV (Thermo Fisher Scientific). Grids were stored under liquid nitrogen until they were imaged.

### Cryo-electron microscopy and tomography

Tilt-series were collected on a Titan Krios 300 kV electron microscope (Thermo Fisher Scientific) equipped with a Gatan K3 direct-electron detector and BioQuantum energy filter (20 eV slit width, Gatan, Pleasanton, CA). A dose symmetric scheme, from - 60° to +60° in 3° increments with groups of two, with a total dose of ∼102 e/Å^2^, using SerialEM.^73,74^ The nominal unbinned pixel size was 1.693 Å and the nominal defocus was −2 to −5 μm in 0.25 µm steps.

Frames were aligned with MotionCor2.^75^ Tilt-series were aligned and processed in IMOD^76^, including CTF correction by phase flipping and dose weighting.^77^ Tomograms were reconstructed in IMOD using weighted back-projection from tilt-series binned by six, four, and two. Bin6 tomograms were processed with IsoNet for denoising and missing wedge correction to enhance visualization of VLPs.^78^ Segmentation of VLPs was done using an EMAN2 convolutional neural network using IsoNet-processed tomograms.^79^ Segmentation was converted to points in EMAN2, and the coordinates were used for initial model files in PEET.^80,81^

### Sub-tomogram averaging of Ty1 VLPs

Initial rounds of alignment and classification were done in PEET 1.15.0. IsoNet-processed tomograms were used to select individual capsomeres as initial references for refinement. All tomograms used for sub-tomogram averaging in PEET were generated using weighted back-projection in 3dmod, with no IsoNet or other processing. Bin 6 particles from six tomograms were aligned to a putative pentamer single particle and separately to a putative hexameric single particle. PCA classification was done after each refinement. Both jobs had classes with hexameric and pentameric capsomeres following classification, the particles were split based on the number of surrounding capsomeres and used in subsequent jobs with a hexameric or pentameric initial model accordingly. The two averages were used as initial references for alignment with an additional eight tomograms, followed by PCA classification. Duplicate particles were removed if they were within 6 pixels (∼51 Å) and 45° of each other. All particles from classes with a hexameric capsomere from these bin6 alignments were combined and aligned at bin4. Particles from classes with a pentameric capsomere were also combined and aligned at bin4. PCA classification was used to remove particles that were not consistent with capsomeres. Another alignment was run at bin4, allowing a full angular search of rotation around the central capsomere with duplicate removal based solely on distance; duplicates within 9 pixels (∼61 Å) were removed. Multiple rounds of PCA classification were then used to classify the particles by both the central capsomere and the organization of surrounding capsomeres. See **Supplemental Table 2** for a description of PEET and RELION jobs. Particles from the selected classes from PEET PCA classifications were then imported into RELION 4.0 for additional alignment and reconstruction at bin 4.^82^

The RELION average of a hexameric capsomere and a pentameric capsomere were then both used separately as initial references for template matching in PEET from the full dataset of 87 tomograms. Initial template matching was done at bin6 in four batches, each containing 23, 23, 23, and 18 tomograms. The resulting averages were subjected to PCA classification, and the retained particles were combined into two batches of 46 and 41 tomograms, which were then aligned at bin 4 in PEET. The particles were subjected to another round of PCA classification. The refined positions and orientations of the retained particles were then imported into RELION 4.0. From sub-tomogram averaging of the L-A virus capsids in the dataset (see below) we determined a calibrated pixel size of 1.645 Å. This pixel size was used throughout the two RELION projects. Particles were aligned at bin 2 with C1 symmetry, followed by 3D classification. Selected particles were then aligned again at bin 2 with C5 symmetry applied for the pentameric capsomere and C2 symmetry applied for the hexameric capsomere. Additional rounds of alignment along with CTF refinement and frame alignment were done as shown in **Supplemental Table 2**. Atomic models were fit into the final structure using UCSF ChimeraX Fit in Map tool.^83^ Q-scores were calculated using the MapQ plugin in UCSF Chimera.^59,83^

### Sub-tomogram averaging of Killer L-A virus

An EMAN2 neural network was trained on manually segmented Killer L-A virus capsids from the bin 6 IsoNet processed tomograms generated as described above. Particle positions were extracted and used for alignment in PEET from two batches of tomograms, each comprising 46 and 41 tomograms. PCA classification was used to remove non-capsid particles, and refined particle positions and orientations were transferred to RELION 4.0 for further processing. Particles were aligned with icosahedral symmetry, as this symmetry was evident from the bin 6 PEET alignment. The final alignment was done at bin2, and the final volume was reconstructed at bin1. A refined pixel size of 1.645 Å was determined based on the fit of PDB 1M1C in the map. This pixel size was used for final refinement and FSC calculation for the L-A virus map.

### Protein expression and purification

The DNA sequence for Ty1 CA-NTD (residues T163-Q260) was synthesized, codon-optimized for expression in *E. coli*, by GeneArt (Thermo Fisher Scientific), amplified by PCR and inserted into a pET22b expression vector (Novagen) between the NdeI and XhoI restriction sites to a produce a C-terminal fusion protein containing the hexa-histidine tag PLEHHHHHH. To obtain ^15^N and ^13^C-^15^N uniformly labelled protein for NMR, Ty1 CA-NTD was expressed in the *E. coli* strain BL21 (DE3) grown in M9 minimal media with ^15^NH_4_Cl or ^15^NH_4_Cl and ^13^C_6_-glucose, as sole nitrogen or nitrogen and carbon sources. Protein expression was induced by addition of 1 mM IPTG to log phase cultures (OD600 = 0.8) followed by incubation overnight at 19 °C with shaking. The cells were harvested, washed with PBS and then resuspended in 50 mM Tris-HCl pH 8.5, 150 mM NaCl, 10 mM Imidazole, 3 mM MgCl_2_ supplemented with 1 mg mL^-1^ lysozyme (Sigma-Aldrich), 10 μg mL-1 DNase I (Sigma-Aldrich) and 1 Protease Inhibitor cocktail tablet (EDTA free, Pierce) per 40 mL of buffer. Cells were lysed using an EmulsiFlex-C5 homogenizer (Avestin) and His-tagged was protein captured from the clarified lysate using immobilized metal ion affinity on a 5 mL Ni^2+^-NTA Superflow column (Qiagen). The column was equilibrated in 50 mM Tris-HCl pH 8.5, 150 mM NaCl, 25 mM Imidazole, 1mM Tris (2-carboxyethyl) phosphine (TCEP) then washed with at least 20 column volumes of the equilibration buffer and Ty1 CA-NTD eluted with 50 mM Tris-HCl pH 8.5, 150 mM NaCl, 250 mM Imidazole. Carboxypeptidase A (Sigma C9268) was added at 1:100 (w:w) ratio and the resulting mixture incubated overnight at 4 °C to digest the C-terminal his-tag. The Carboxypeptidase A was then inactivated by the addition of 5 mM TCEP and Ty1 CA-NTD further purified by gel filtration chromatography on a Superdex 75 (26/60) column equilibrated in 20 mM Tris-HCl, 150 mM NaCl, 1mM TCEP pH 7.5. Ty1 CA-NTD containing fractions were concentrated to 1–2 mM by centrifugal ultrafiltration (Vivaspin, MWCO 10 kDa), then snap frozen and stored at -80 °C. Protein concentration was determined by UV absorbance spectroscopy using an extinction coefficient calculated at 280 nm derived from the tryptophan and tyrosine content.

### NMR spectroscopy

NMR experiments were performed on samples ranging between 200 to 600 µM of uniformly [^15^N,^13^C]-labelled Ty1 CA-NTD in 50 mM phosphate buffer at pH 5, 100 mM NaCl, 1 mM TCEP with 5% D_2_O, and 0.01% NaN_3_. Experiments were carried out at 25 °C. Backbone assignments were obtained using the standard set of triple resonance experiments^84^ including HNCO, HN(CA)CO, HNCA, HN(CO)CA, HNCACB, CBCA(CO)NH, HCA(CO)N, HCAN and (H)N(CA)NNH from data collected on Bruker spectrometers operating at fields between 600–950 MHz. Resonances corresponding to the aliphatic sidechains were assigned using a combination of H(C)CH-TOCSY, (H)CCH-TOCSY and amide-detected (H)C(CCO)NH- and H(CCCO)NH-TOCSY spectra. Resonances for the aromatic sidechains were obtained using a 3-dimensional ^13^C-NOESY-HSQC in conjunction with the H(C)CH-TOCSY, (H)CCH-TOCSY optimized for aromatic resonances; the sidechain amino positions of Asn and Gln residues resonances were assigned using a 3-dimensional ^15^N-NOESY-HSQC. Distance restraints were derived from the 3-dimensional ^15^N-NOESY-HSQC and 3-dimensional ^13^C-NOESY-HSQC experiments together with a 3-dimensional ^13^C-NOESY-HSQC (optimized for aliphatic resonances). All NOESY experiments employed mixing times of 120 milliseconds.

Residual dipolar couplings (RDCs) data were collected using [^13^C,^15^N]-Ty1 CA-NTD on a 300 µM sample in the presence and absence of 14 mg.mL^-1^ of Pf1 phage. Measurements of ^1^D_NH_ RDCs were conducted using the TROSY-based ARTSY pulse sequence^85^ on an 800 MHz Bruker spectrometer. The ^1^D_NH_ RDCs range from approximately + 20 to -14 Hz. Data were processed using NMRPipe^86^ and analysed in CARA.^87^

### NMR structure determination

The family of 20 NMR structures of Ty1 CA-NTD was calculated using conformational restraints for 2,251 inter-proton distances and 72 backbone dihedral angles (**Supplemental Table 3**). The set of backbone φ (phi) and ψ (psi) dihedral angles was obtained from the analysis of the ^1^HN, ^1^Hα, ^15^N, ^13^Cα, ^13^Cβ and ^13^C′ chemical shifts using the TALOS-N software^88^ for the residues located in the well-ordered regions of the protein core, as defined by NMR relaxation experiments and the RCI (Random Coil Index) approach.^89^ NOEs, used as distance restraints in structure calculation, were obtained from the analysis of cross-peak intensities in the 3D ^13^C-^1^H and ^15^N-^1^H HSQC-NOESY spectra. The intra-residue and sequential cross-peaks of NOESY spectra were assigned manually, while the remainder of the cross-peaks were assigned using the automatic iterative procedure of spectra assignment/structure calculation implemented in ARIA 2.3 software^90^ using the ARIA web-server ^91^. The automatic assignment and the inter-proton distances provided at the last iteration of the ARIA 2.3 protocol were further manually verified by multiple steps of the structure refinement accomplished using the simulated annealing protocol of the CNS 1.21 software package.^92^ The RDC values for regions of defined secondary structure were introduced using the SANI option starting from the 5^th^ ARIA iteration. The required rhombicity and tensor magnitude values were derived independently using Module^93^ from a previous round of structure calculation that did not use RDC restraints. Structure refinement included a high-temperature torsion-angle molecular dynamics stage followed by a slow-cooling torsion-angle phase, a second slow-cooling phase in Cartesian space and Powell energy minimization. Structure refinement was performed until no NOE violations larger than 0.5 Å and no dihedral angle violations greater than 5° occurred. In the final iteration of the refinement protocol 40 structures were calculated. From these, a family of 20 NMR structures was selected in accordance with the lowest-energy criterion. The statistics for NMR structure determination are presented in **Supplemental Table 3**. Structure visualization and analysis were carried out using PyMOL 2.6.2 (Schrödinger LLC). Structure quality was assessed using the PSVS web-server^94^ and deposited in the PDB with accession code 9TGB.

### NMR protein dynamics analysis

Ty1 CA-NTD dynamics were examined by measuring backbone ^15^N relaxation parameters; longitudinal relaxation rate (*R*_1_), transverse relaxation rate (*R*_2_) and heteronuclear [^1^H]-^15^N-NOE (HetNOE)^95^ at 298 K and ^1^H spectrometer frequencies of 600 and 950 MHz. *R*_1_ and *R*_2_, were determined by acquiring 2D heteronuclear ^1^H-^15^N-HSQC spectra with varying relaxation period varied. For *R*_1_ measurements delay times of 10, 100, 200, 400(2), 500, 750 , 1000, 1250, and 1500 ms were applied. For *R*_2_ the delay times were 8, 16(2), 32, 48, 64(2), 80, 96, 128, 160, and 192 ms. The resonance intensities at each relaxation delay were quantified using NMRPipe and R_1_ and R_2_ obtained by fitting individual peak intensities using nonlinear spectral lineshape modelling to a single exponential using routines also within NMRPipe. HetNOE values were calculated from peak intensity ratios obtained from interleaved ^1^H-^15^N spectra, 5 s delay, recorded with and without ^1^H saturation prior to the ^15^N excitation pulse. The average errors were estimated at 3% for the R_1_ and R_2_ measurements and at 5% on the steady-state HetNOE values.

### ^15^N-Carr-Purcell-Meiboom-Gill (CPMG) relaxation dispersion measurements

^15^N CPMG relaxation dispersion experiments were recorded at 600 MHz (14.1 T) and 950 MHz (22.3 T). An evolution CPMG delay (T-_CPMG_) of 40 ms was employed with CPMG frequencies (*ν_CPMG_*) of 25, 50, 100(2), 200, 400(2), 700 and 1000, Hz. An experiment recorded without CPMG delay served as a reference for the calculation of R_2,eff_ rates as a function of CPMG field as described.^96^ Uncertainties in R_2,eff_ values were obtained from duplicate measurements at two different ν_CPMG_ frequencies. Spectra were processed and peak intensities measured in NMRPipe with overlapping peaks excluded from further analysis. Peak intensities were used to calculate the effective transverse relaxation rates using R_eff2_=−1/T ln(I_νCPMG_/I_0_), where *T* is the constant CPMG time, and *I_νCPMG_* and *I*_0_ are the signal intensities in the presence or absence of the CPMG pulse, respectively. Fitting of relaxation dispersion experiments at the two fields was performed simultaneously with NESSY^97^ with data fitted to no- (model-1), fast- (Model-2) (Wang et al., 2001) and slow-(Model-3)^56^ exchange models. Selection of the best model was undertaken using Akaike information criteria^98^ with parameters were held constant for each residue, except for *δω* (model 3) and Φ (model 2), which were weighted by the heteronuclear frequency (^15^N). Errors in extracted parameters were estimated using 500 Monte Carlo simulations implemented in NESSY.

### Structural similarity searches

The PDBeFold (https://www.ebi.ac.uk/msd-srv/ssm/) comparison server was used to perform similarity searches and align structural homologues of Ty1 CA-NTD.

## Data Availability

The cryo-ET volumes from sub-tomogram averaging in this study have been deposited in the Electron Microscopy Data Bank (www.emdatabank.org) under the following accession numbers: TBD. The co-ordinates for Ty1 CA-NTD structure have been deposited in the Protein Data Bank with accession number 9TGB and chemical shift assignments in the BioMagResBank accession number 53457. All relevant data are available from the corresponding author upon request.

## Acknowledgements

This work was supported in part by the University of Wisconsin, Madison, the Department of Biochemistry at the University of Wisconsin, Madison, and public health service grants R01 GM114561 and U24 GM139168 to E.R.W. from the NIH. This work was also funded by NIH grants to DJG (R01GM124216 and R01GM156837) and by the Francis Crick Institute, which receives its core funding for IAT from Cancer Research UK (CC2029), the UK Medical Research Council (CC2029) and the Wellcome Trust (CC2029). We are grateful for the use of facilities and instrumentation at the Cryo-EM Research Center in the Department of Biochemistry at the University of Wisconsin, Madison and the UK biomedical NMR facility at the Francis Crick Institute. Molecular graphics and analyses performed in part with UCSF Chimera, developed by the Resource for Biocomputing, Visualization, and Informatics at the University of California, San Francisco, with support from NIH P41-GM103311. We are grateful for the computational resources supplied through the SBGrid Consortium (Morin et al., 2013).

## Author Contributions

Bryan S. Sibert: Conceptualization, Formal Analysis, Investigation, Validation, Visualization, Writing – Original Draft Preparation, Writing – Review & Editing.

J. Adam Hannon-Hatfield: Conceptualization, Formal Analysis, Investigation, Validation, Visualization, Writing – Original Draft Preparation, Writing – Review & Editing.

Giuseppe Nicastro: Formal Analysis, Investigation, Validation, Visualization, Writing – Review & Editing.

Matthew A Cottee: Formal analysis, Investigation, Validation, Visualization.

Ian A. Taylor: Conceptualization, Formal Analysis, Funding Acquisition, Supervision, Investigation, Visualization, Writing – Review & Editing.

David J. Garfinkel: Conceptualization, Funding Acquisition, Supervision, Writing – Original Draft Preparation, Writing – Review & Editing.

Elizabeth R. Wright: Conceptualization, Visualization, Funding Acquisition, Supervision, Writing – Original Draft Preparation, Writing – Review & Editing.

## Competing Interests

The authors declare no competing financial or non-financial interests.

**Correspondence and requests for materials** should be addressed to D.J.G. or E. R. W. or I. A. T.

## Supplemental Figures and Legends, Supplemental Tables, and Supplemental Movie Legends

**Supplemental Figure 1.**
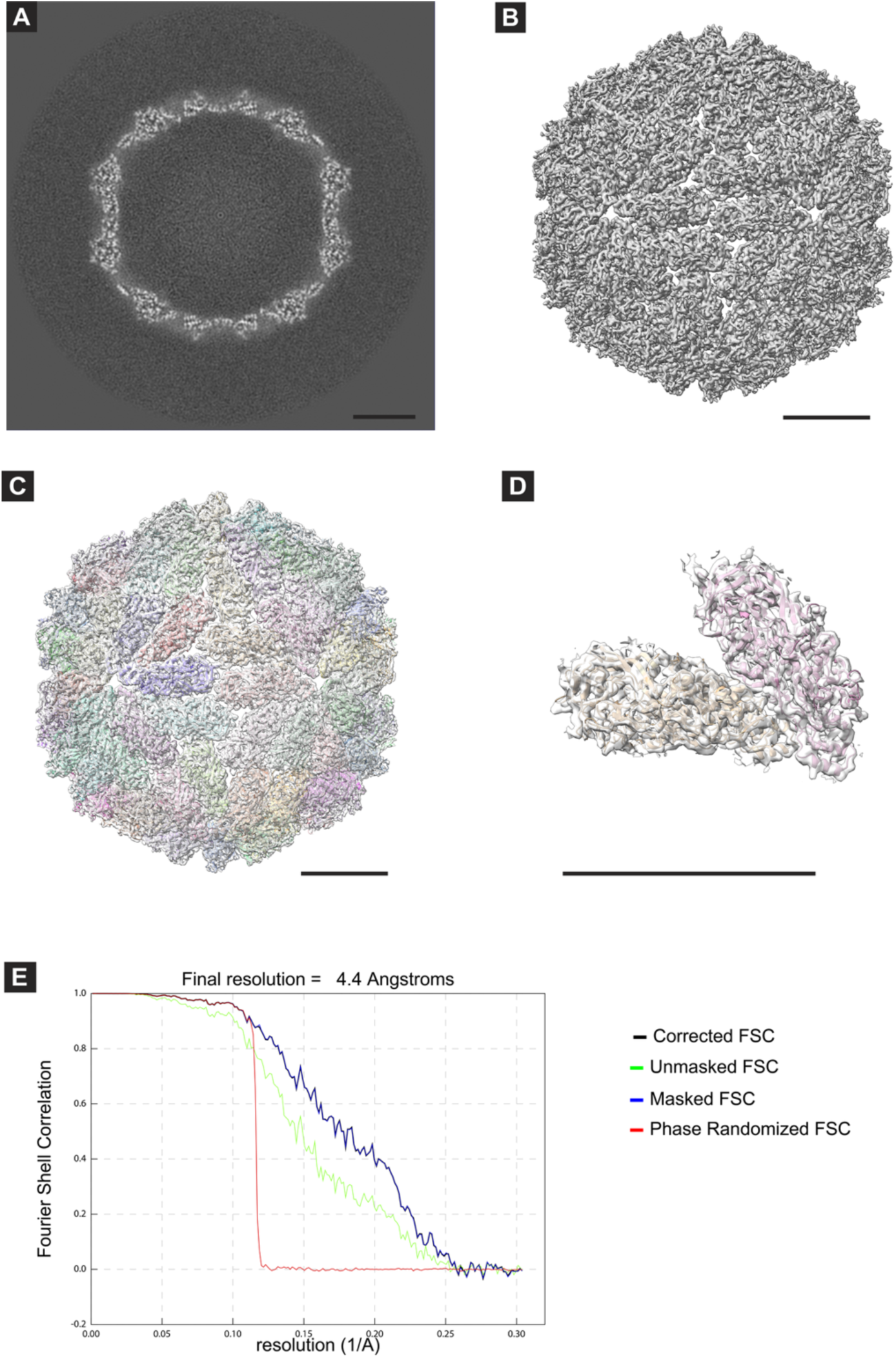
Sub-tomogram average of LA virus capsid. (A) Single X, Y slice from the electron density map from sub-tomogram averaging of an LA virus capsid. (B) Isosurface of the map in A. (C) Atomic models from PDB 1M1C fit into the isosurface in B. (D) The asymmetric unit of the LA virus capsid, comprised of two copies of the monomer, is shown with surrounding density from the subtomogram average. 100 nm scale bars in A, B, C, D. (E) FSC plot generated by Relion. Red line - phase randomized maps, green line - unmasked maps, blue line - masked maps, black line - corrected, masked maps.

**Supplemental Figure 2.**
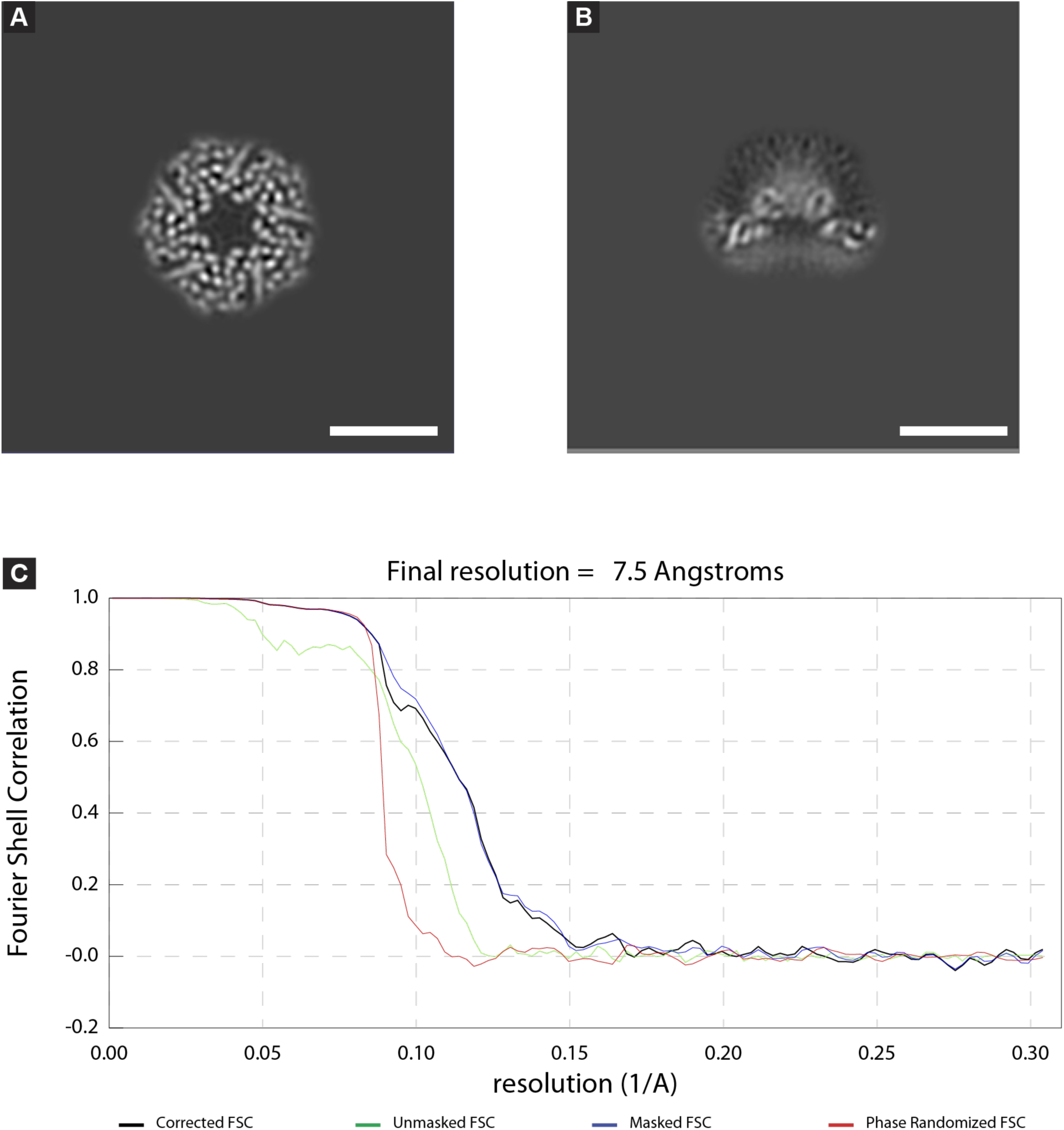
FSC and local resolution of Ty1 VLP pentameric capsomere. (A) Masked single X, Y slice from electron density map from sub-tomogram averaging of a pentameric capsomere from Ty1 VLPs. (B) Masked single X, Z slice from the map shown in A. (C) FSC plot generated from the two half-maps in Relion. Black - Masked, corrected FSC plot, Green - Unmasked, Blue - Masked, Red - Phase randomized. Scale bars are 10 nm.

**Supplemental Figure 3.**
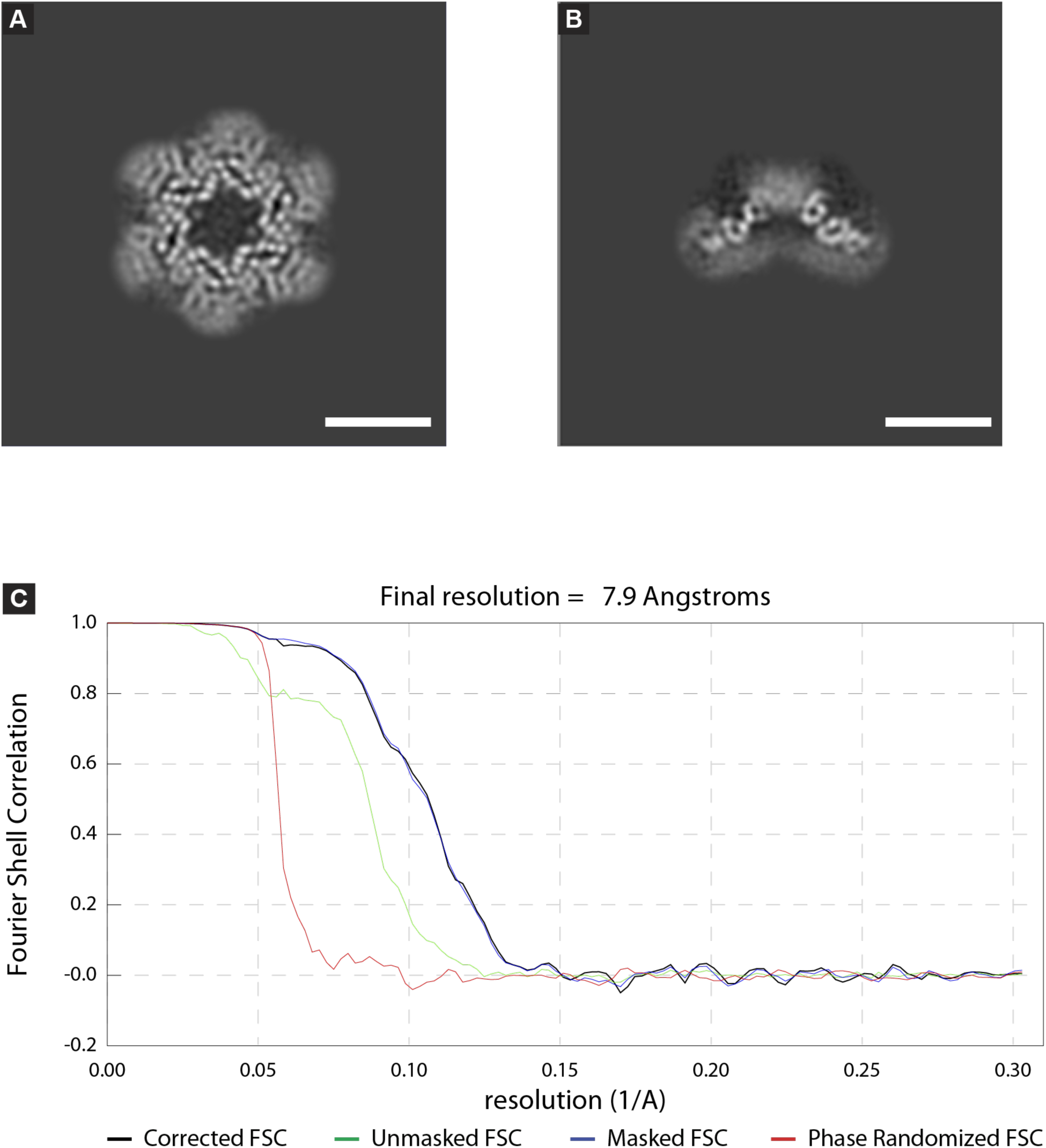
FSC and local resolution of Ty1 VLP hexameric capsomere. (A) Masked single X, Y slice from electron density map from sub-tomogram averaging of a hexameric capsomere from Ty1 VLPs. (B) Masked single X, Z slice from the map shown in A. (C) FSC plot generated from the two half-maps in Relion. Black - Masked, corrected FSC plot, Green - Unmasked, Blue - Masked, Red - Phase randomized. Scale bars are 10 nm.

**Supplemental Figure 4.**
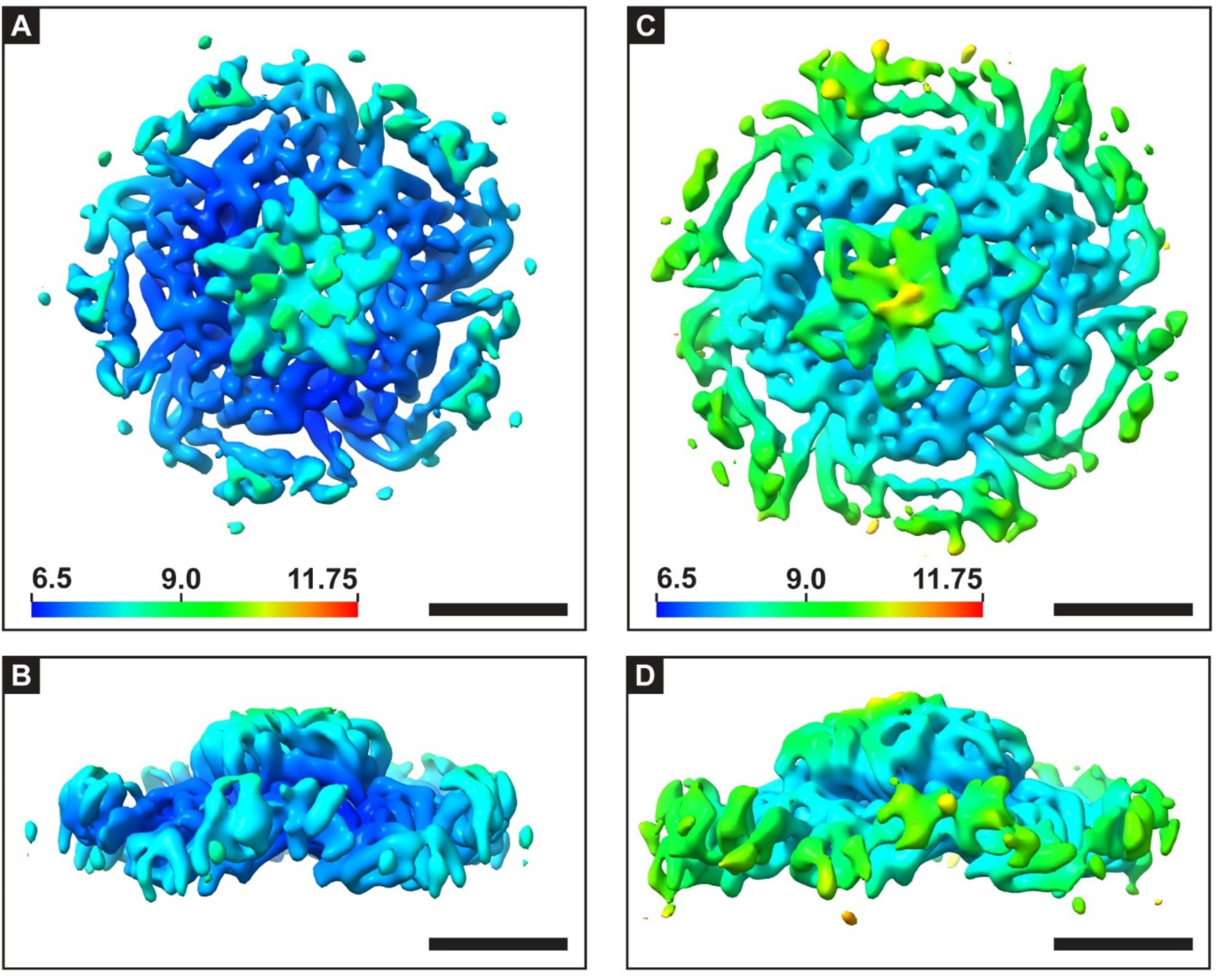
Local resolution maps of Ty1 VLP capsomeres. (A) Isosurface of Ty1 pentameric capsomere colored by local resolution as indicated by the color key in the lower left (units in Å). Minimum and maximum resolutions at the displayed threshold are 6.6 Å and 8.7 Å, respectively. (B) Isosurface in A rotated 90° around the x-axis. (C) Isosurface of Ty1 hexameric capsomere colored by local resolution as indicated by the color key in the lower left (units in Å). Minimum and maximum resolutions at the displayed threshold are 7.2 Å and 11.4 Å, respectively. (D) isosurface in C rotated 90° around the x-axis. Scale bar 5 nm

**Supplemental Figure 5.**
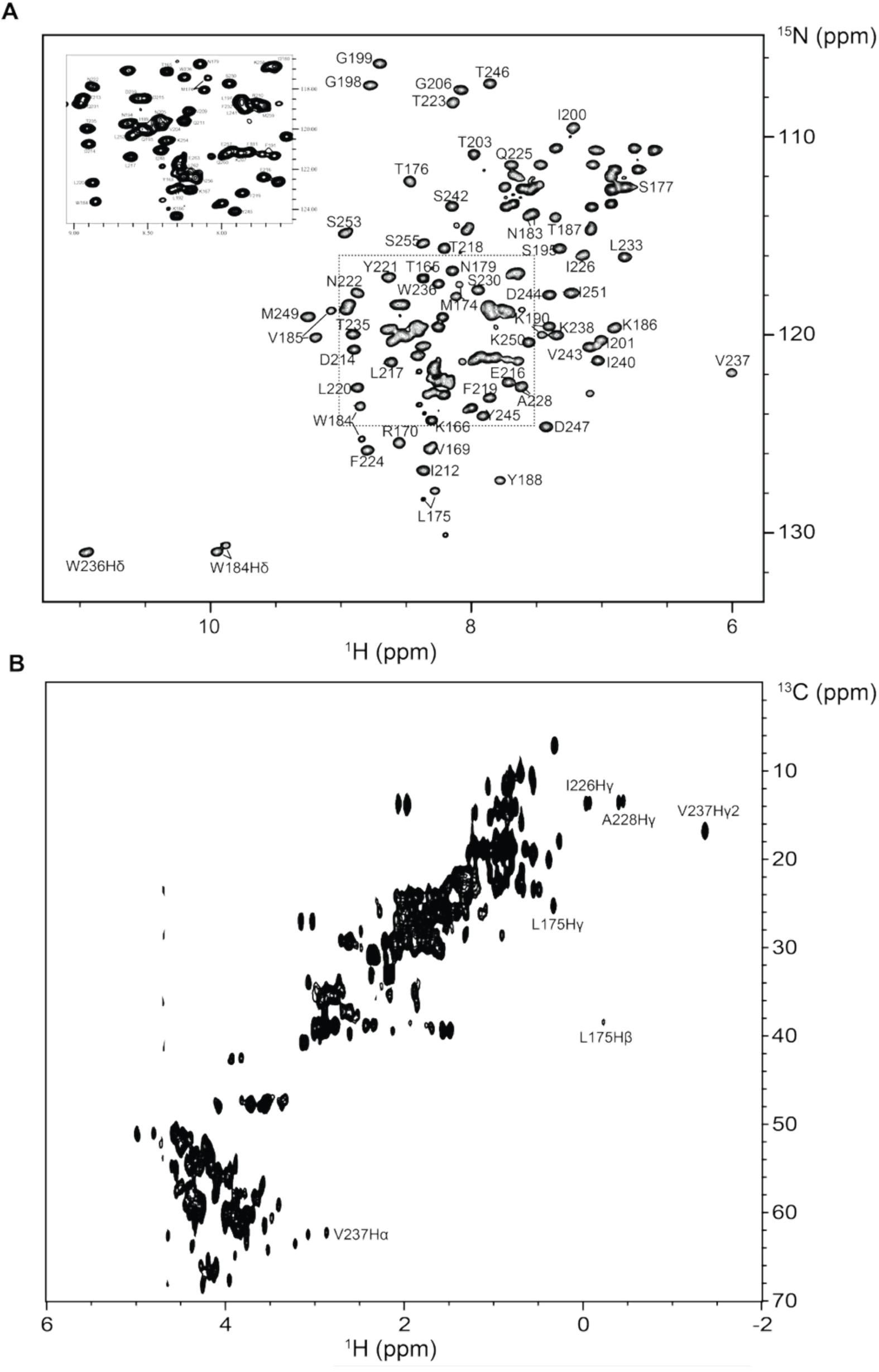
TY1 CA-NTD NMR spectra. (A) Assigned ^1^H-^15^N HSQC spectrum of uniformly ^13^C-^15^N labelled TY1-CA-NTD. Backbone amide resonances are labelled along with side-chain tryptophan NH. Inset is an expanded view of the boxed central region of the spectrum. Cross peaks corresponding to residues M174, L175, N183, W184, V185, K190 and F191 that have two resonances resulting from major and minor conformations in slow exchange are indicated with lines (B) ^1^H-^13^C HMQC spectrum of uniformly ^13^C-^15^N labelled TY1-CA-NTD. Selected residues that show a high degree of perturbation through proximity-induced ring current effects are labelled.

**Supplemental Figure 6.**
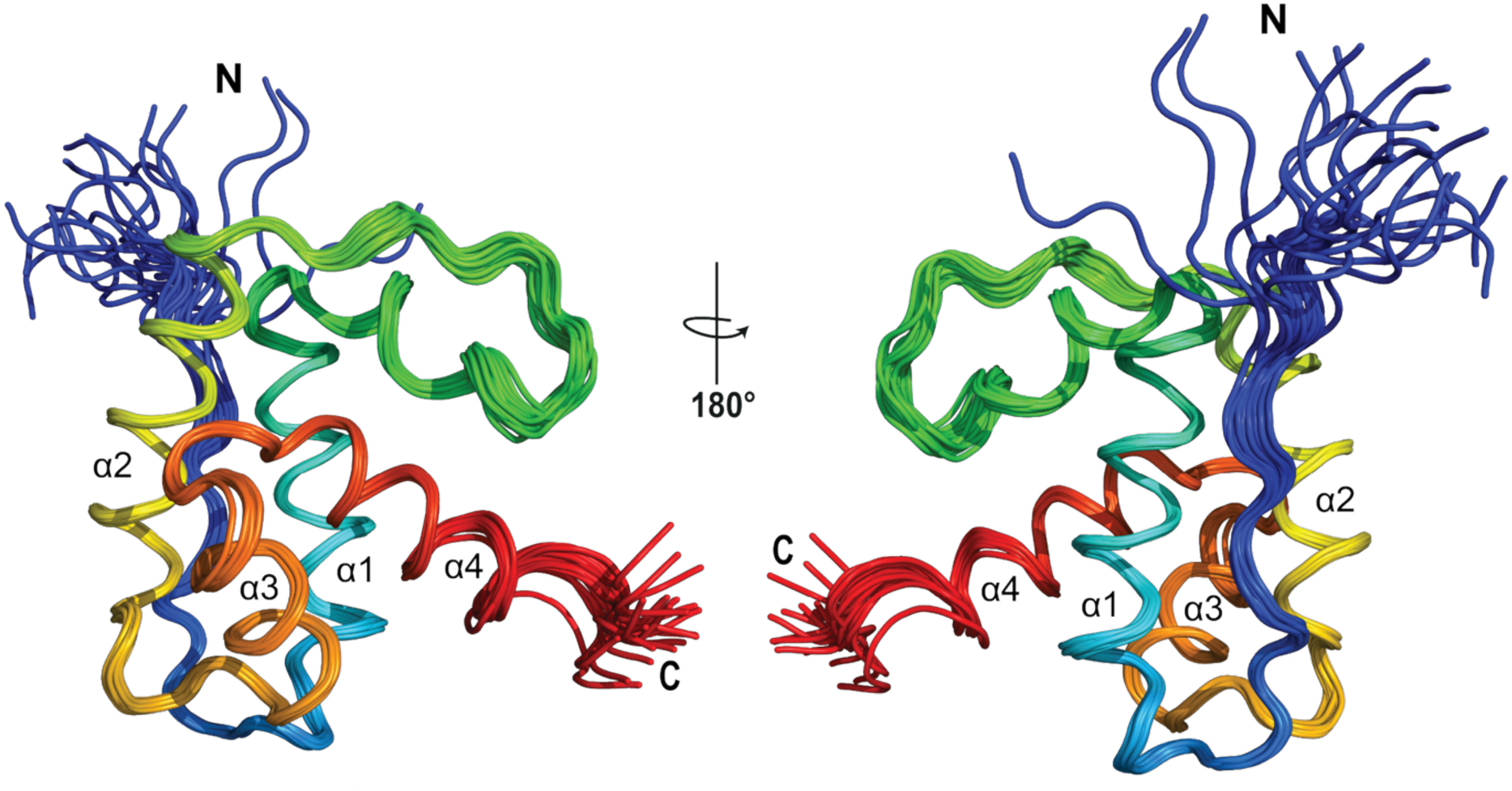
Family of TY1 CA-NTD solution NMR structures. The protein backbone for each of the 20 conformers in the final refinement is shown in ribbon representation in two views rotated by 180 °. The backbone is colored from the N-to C-terminus in blue to red with α-helices labelled sequentially.

**Supplemental Figure 7.**
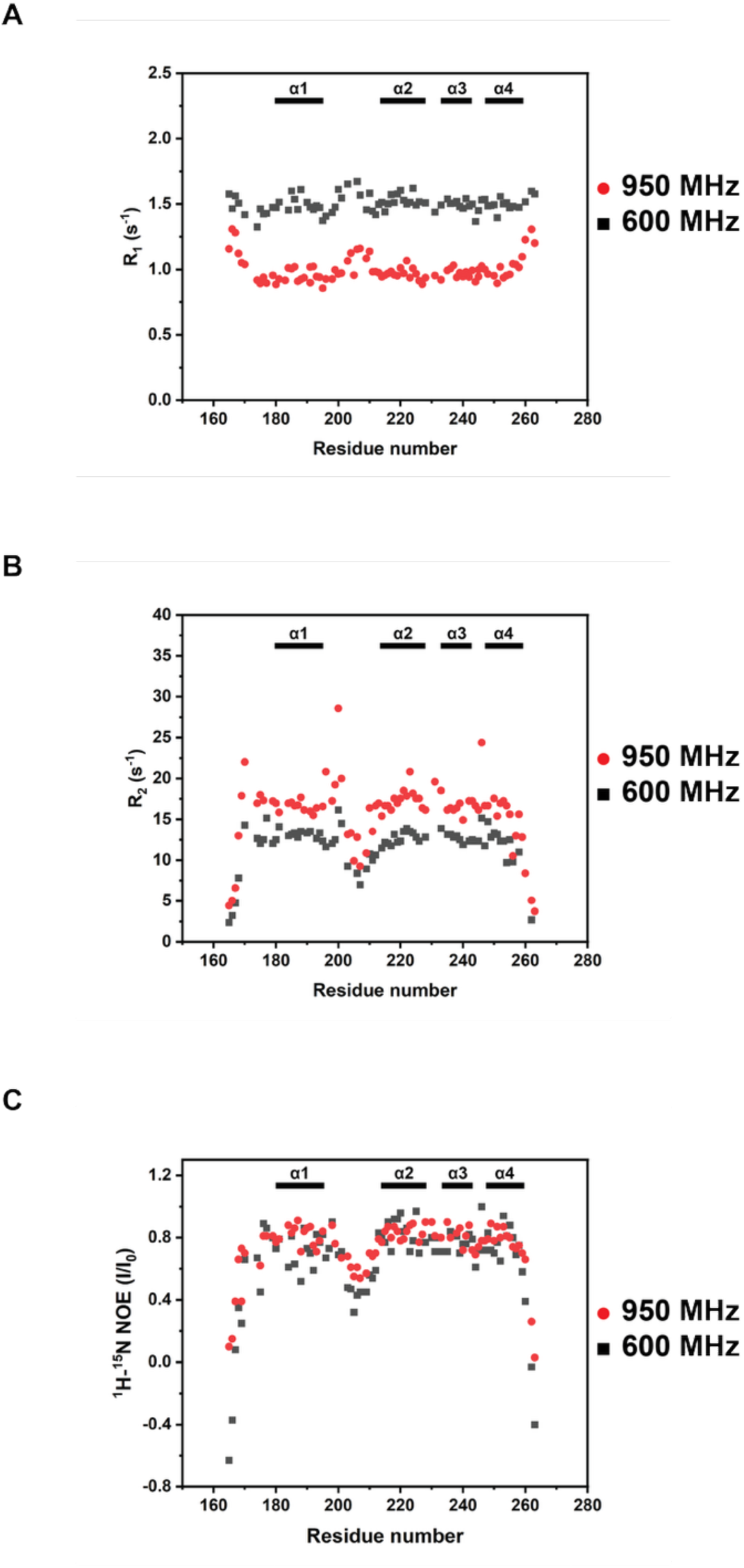
NMR Relaxation data. The Backbone ^15^N relaxation parameters of TY1 CA-NTD recorded at 600 MHz and 950 MHz at 25 °C. (A) the spin-lattice relaxation rate R_1_, (B) the spin-spin relaxation rate R_2_ and (C) the steady-state heteronuclear ^1^H-^15^N NOE recorded for each residue is plotted against sequence position. The bars in each plot represent the positions of α-helices in the protein. In each plot, residues in the long α1-α2 connecting loop show increased fast motions relative to the core domain.

**Supplemental Figure 8.**
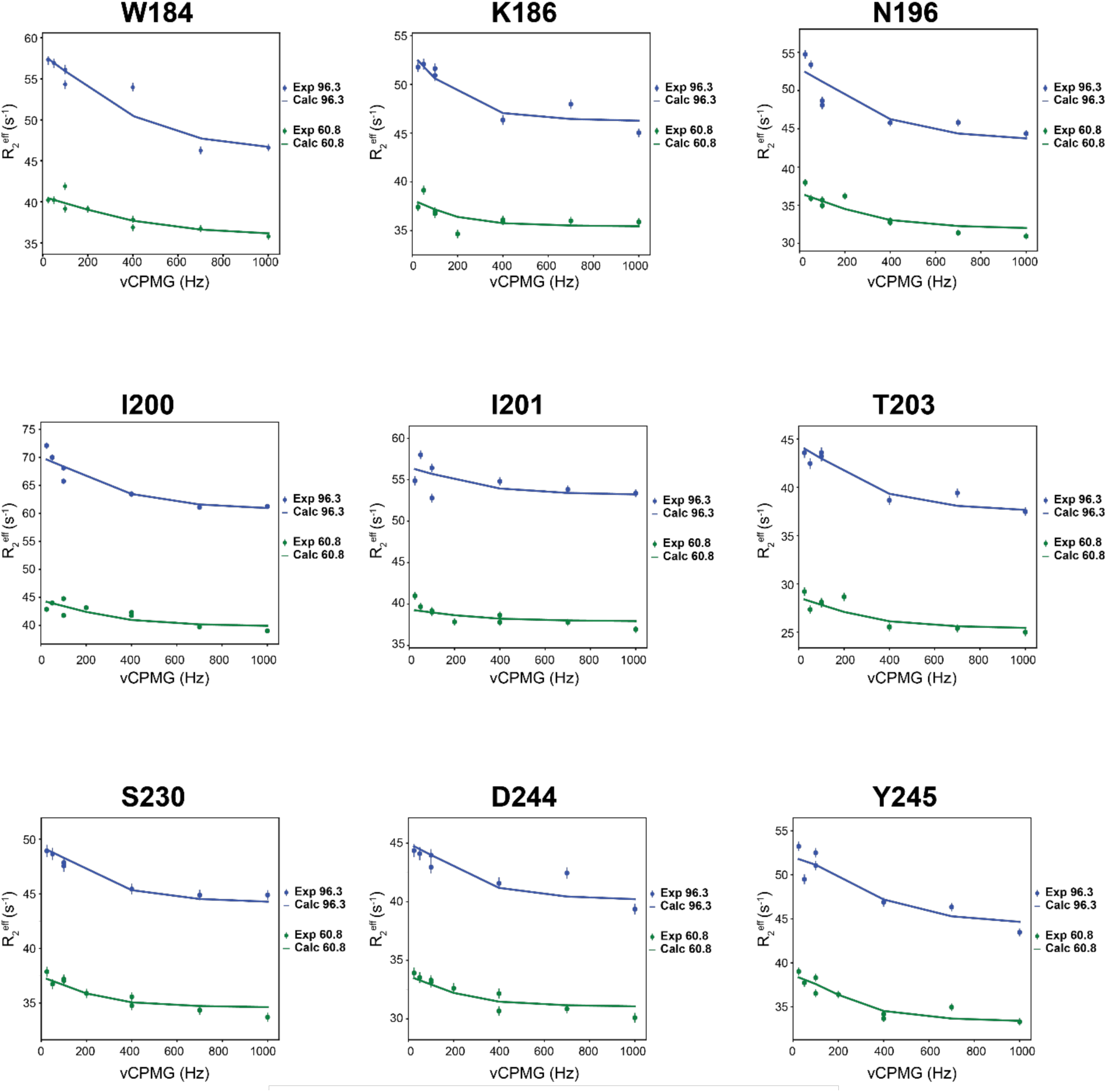
^15^N R_2_^eff^ relaxation dispersion profiles. Representative plots of R_2_^eff^ against vCPMG are shown for assignable residues that show exchange. Data were recorded at 600 MHz (green) and 950 MHz (blue) ^1^H frequency; the legend indicates the corresponding^15^N frequency. The solid lines indicate the best fit to a global two-site slow exchange model to derive P_b_ and R_ex_ parameters. Errors were assessed using 500 steps of Monte Carlo simulation.

**Supplemental Figure 9.**
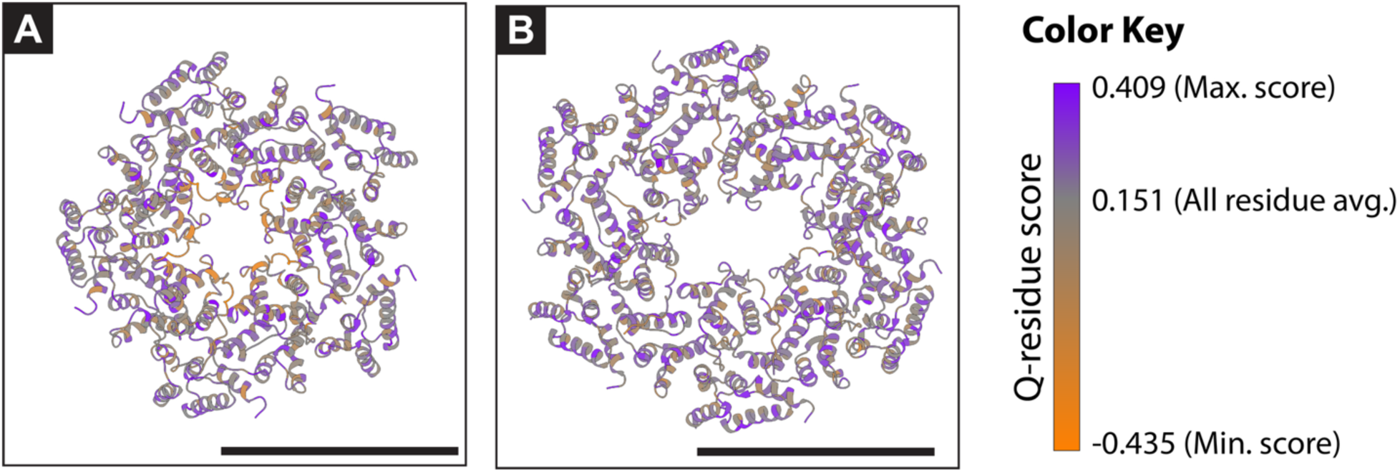
Atomic models fit into density maps colored by Q-score. (A) Five copies of CA-CTD Dimer-1 (PDB ID:7NLH) comprising ten CA-CTD monomers and five copies of the CA-NTD NMR structure fit into the pentameric STA density map. Residues are colored by Q-residue score, as indicated by the color key to the right. The expected Q-score with a map resolution of 7.5 Å is 0.129; the average Q-score for the CA-NTD residues 163-263 is 0.103. The average Q-score for the CA-CTD residues 261-351 is 0.165. (B) As in A for the hexameric STA density map. The expected Q-score with a map resolution of 7.9 Å is 0.123; the average Q-score for the CA-NTD residues 163-263 is 0.145. The average Q-score for the CA-CTD residues 261-351 is 0.165. Scale bars are 10 nm.

**Supplemental Figure 10.**
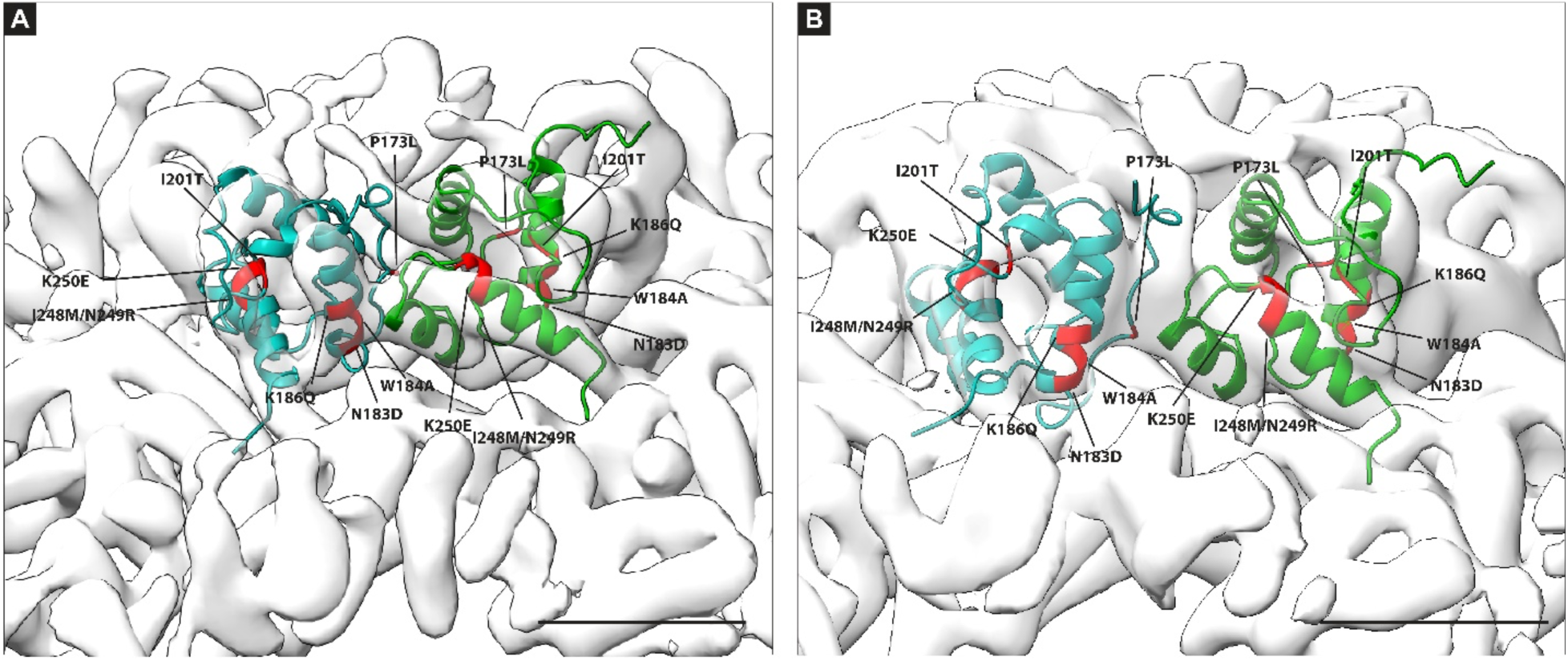
CA-NTD mutations with known phenotypes. (A) Sub-tomogram average of a pentameric capsomere with 2 copies of the CA-NTD structure fitted into the central turret shown in green and teal cartoon as in Figure 6. Position of previously characterized mutations in the CA-NTD are indicated. (B) As in A for the hexameric capsomere. Scale bars are 10 nm.

**Supplemental Figure 11.**
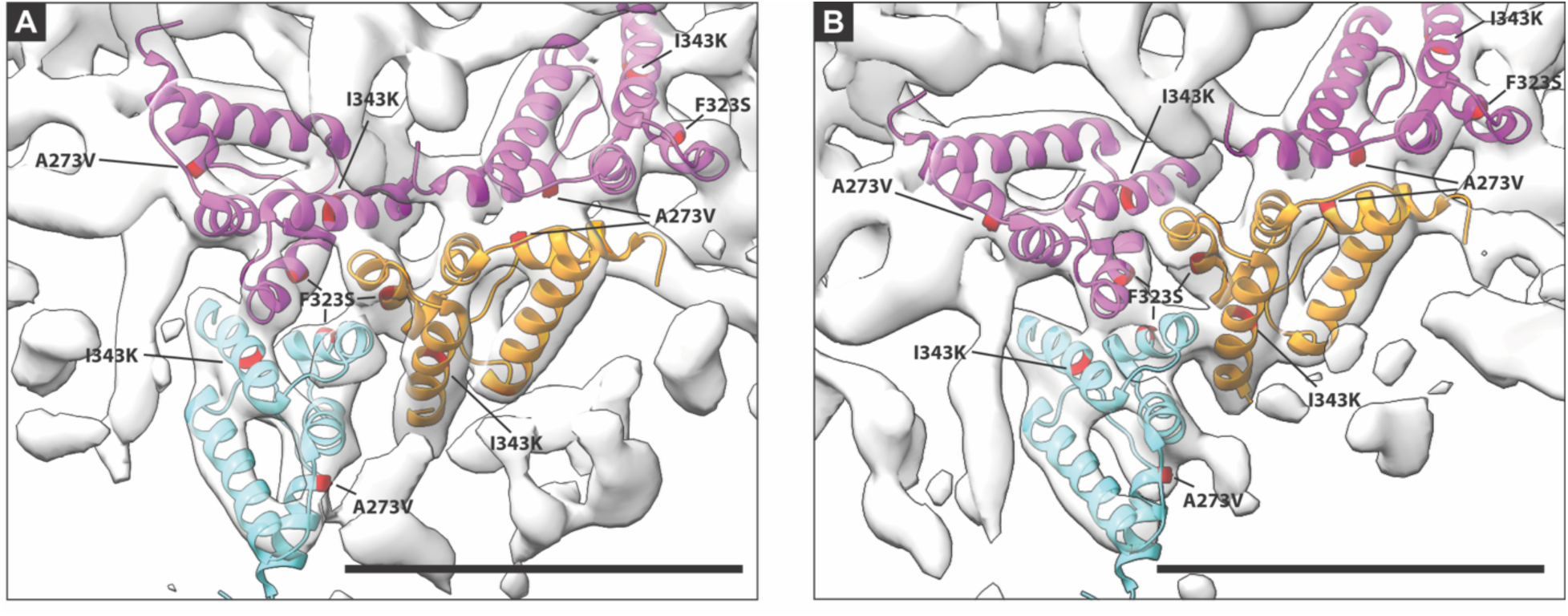
CA-CTD mutations with known phenotypes. (A) Sub-tomogram average of a pentameric capsomere with models from p18m (PDB ID: 7NLH) fitted into the map density. Four copies of the Ty1 CA-CTD are shown including dimeric chains A (magenta) and B (tangerine) surrounding the central capsomere and an additional copy of chain A (cyan) shown fit into neighboring capsomere density. Previously characterized mutations within CA-CTD are indicated. (B) As in A for the hexameric capsomere. Scale bars are 5 nm.

**Table S1.** Data collection parameters for cryo-ET.

|  |  |
| --- | --- |
| **Microscope** | TFS Titan Krios G3i |
| **Voltage (kV)** | 300 |
| **Energy filter** | Gatan Bioquantum, 20eV slit |
| **Detector** | Gatan K3 |
| **Detector mode** | CDS counting mode, super resolution |
| **Frames per tilt** | 8 |
| **Magnification** | 53000x |
| **Total e<sup>-</sup> dose** | 102 e<sup>-</sup>/Å<sup>2</sup> |
| **Nominal defocus** | 2.5 - 5 μm |
| **Pixel size (nominal)** | 1.693 Å (non-super-res) |
| **Pixel size (calibrated)** | 1.645 Å (non-super-res) |
| **Software** | SerialEM |
| **Tilt-scheme** | Dose symmetric, -60/+60°, 3° increments, grouped by 2 |
| **Tomograms included** | 87 |

**Table S3.**
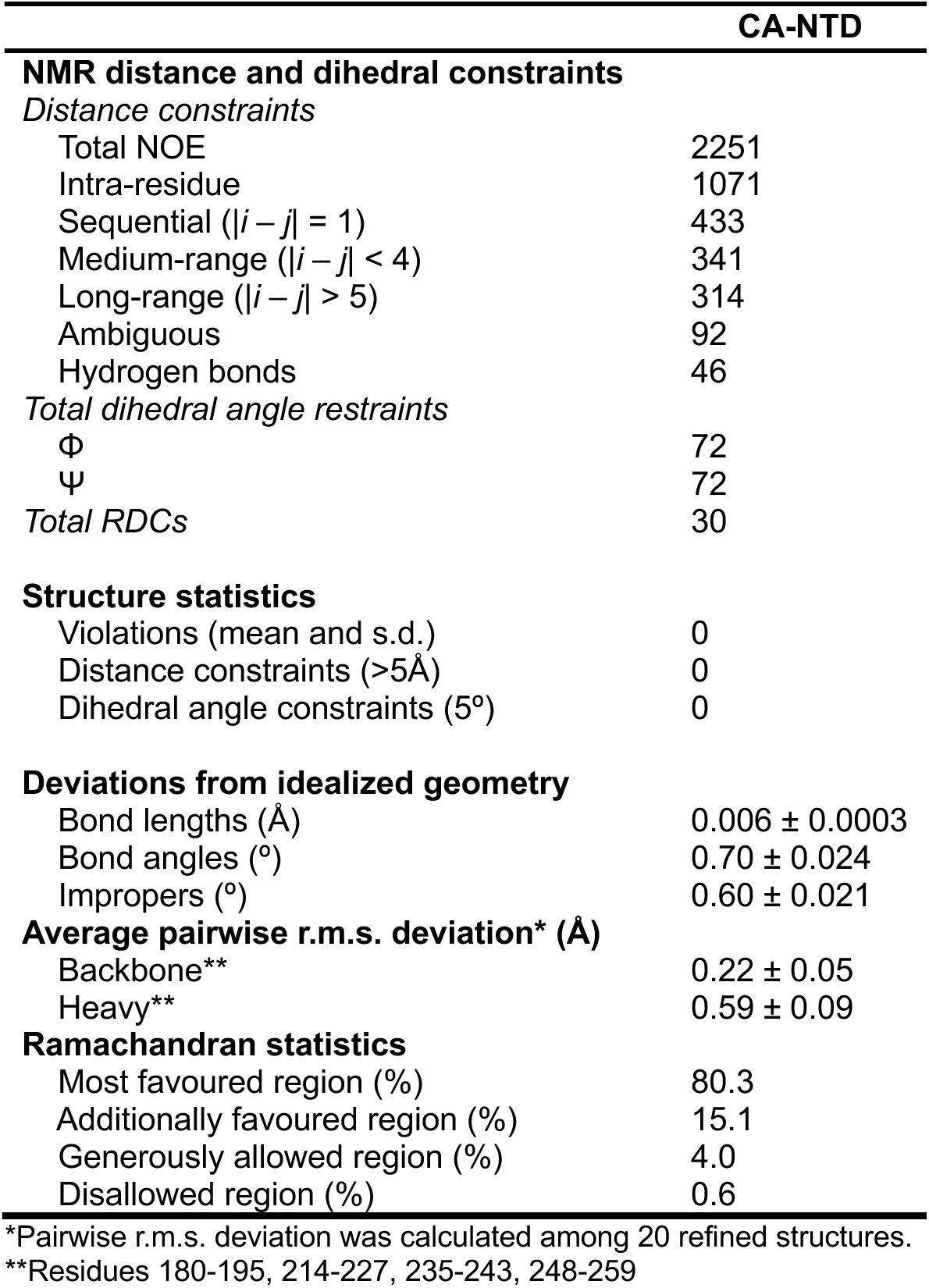
NMR and refinement statistics.

**Table S4.**
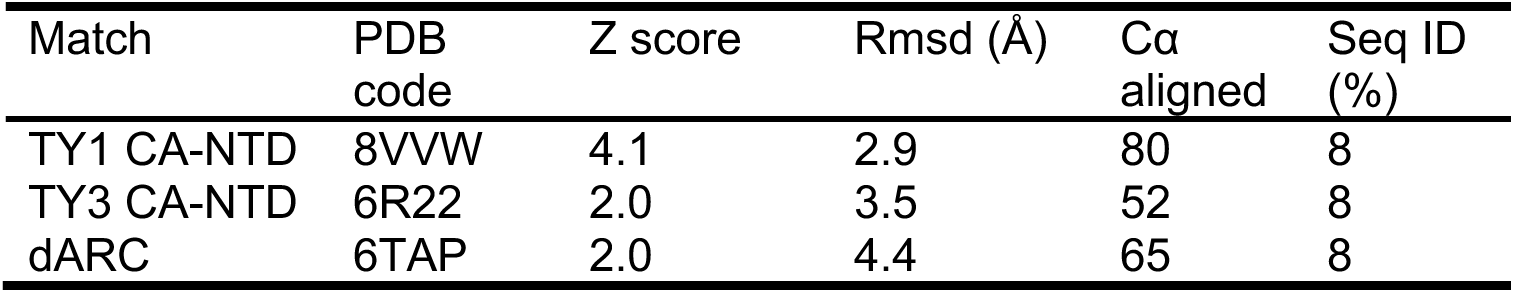
PDB structural similarity search.

**Supplemental Movie 1. Density map and atomic model fitting into the pentameric Ty1 VLP capsomere.** An isosurface of the density map is displayed as viewed from outside of the virion to help orient the viewer. This is followed by the display of X, Y slices of the density map moving through Z from the outside of the virion towards the interior, away from the viewer. The isosurface is shown again and rotated 90° around the X axis, immediately followed by the display of Y, Z slices of the density map moving through the volume, away from the viewer. The isosurface is displayed again, with atomic models of the CA-CTD (PDB:7NLH, chain A magenta, chain B tangerine) and the CA-NTD (NMR structure, green) fit in the density. The isosurface is rotated around the X and Y axes to show the structure and fit from several angles. The isosurface is then removed to show the organization of the fit atomic models more clearly.

**Supplemental Movie 2. Density map and atomic model fitting into the hexameric Ty1 VLP capsomere.** An isosurface of the density map is displayed, viewed from outside the virion, to help orient the viewer. This is followed by the display of X, Y slices of the density map moving through Z from the outside of the virion towards the interior, away from the viewer. The isosurface is shown again and rotated 90° around the X axis, immediately followed by the display of Y, Z slices of the density map moving through the volume, away from the viewer. The isosurface is displayed again and shown with atomic models of the CA-CTD (PDB:7NLH, chain A magenta, chain B tangerine) and CA-NTD (NMR structure, green) fit in the density. The isosurface is rotated around the X and Y axes to show the structure and fit from several angles. The isosurface is then removed to show the organization of the fit atomic models more clearly.

